# Comprehensive investigation of SARS-CoV-2 intestinal pathogenesis in *Drosophila*

**DOI:** 10.1101/2025.06.25.661044

**Authors:** Layla El Kamali, Peter Nagy, Justine Girard, Nicolas Buchon, Patrick Mavingui, Chaker El-Kalamouni, Dani Osman

**Author notes:** Correspondence (C.EK.); (D.O.).

## Abstract

Gastrointestinal (GI) manifestations have been increasingly reported in COVID-19 patients. Here, we use the *Drosophila melanogaster* midgut model to investigate SARS-CoV-2-induced GI pathogenesis. The fly midgut exhibits susceptibility to orally administered virus, resulting in disrupted epithelial architecture, reduced organ size, and altered visceral muscle dynamics. These effects are accompanied by sustained proliferation of intestinal stem cells alongside decreased replenishment and viability of differentiated cells. Transcriptomic profiling reveals biphasic perturbations in midgut gene expression, particularly in pathways related to lipid metabolism. Intriguingly, SARS-CoV-2 elicits a dichotomous effect on lipid homeostasis, with lipid droplet accumulation in the posterior midgut and depletion in anterior segments. Treatment with Plitidepsin, a COVID-19 drug candidate, mitigates most SARS-CoV-2 pathogenic features in both the *Drosophila* midgut and human pulmonary cells, while modulating basal lipid droplet homeostasis in uninfected conditions. These findings establish the *Drosophila* midgut as a potent model for studying SARS-CoV-2 GI pathogenesis and evaluating antiviral compounds.

## Introduction

Coronaviruses (CoVs), a group of enveloped positive-sense single-stranded RNA viruses, predominantly affect birds and mammals, causing a spectrum of diseases ranging from mild to fatal [1,2]. These viruses target predominantly the epithelial cells of the gastrointestinal (GI) and airway tracts, causing gastroenteritis and respiratory illnesses across various animal species, including wild, domestic, and farmed populations [3]. Certain CoVs have breached the interspecies barrier, leading to zoonotic diseases, with the severe acute respiratory syndrome coronavirus 2 (SARS-CoV-2) being a prime example [4]. This virus, belonging to the β-coronavirus genus, has been identified as the causative agent of the global pandemic known as coronavirus disease 2019 (COVID-19).

While respiratory complications are the primary clinical concern in SARS-CoV-2 infections, emerging evidence indicates the virus’s potential to disrupt multiple organ systems, particularly the digestive system [5]. Cohort studies have documented a significant prevalence of GI symptoms, such as diarrhea, vomiting, loss of appetite, and abdominal discomfort, in COVID-19 patients, with occurrences ranging from 12 to 61% (reviewed in [5]). Intriguingly, these symptoms can manifest before or even in the absence of respiratory signs [6], suggesting a direct viral invasion of the GI tract as an alternative infection route. This is further supported by the expression of the angiotensin-converting enzyme 2 (ACE2) receptor, which acts as the initial cell-virus contact point, and the transmembrane serine protease 2 (TMPRSS2), which primes the viral spike protein for entry, in GI epithelial tissues [7–9]. This hypothesis is bolstered by the prolonged detection of viral antigens and RNA in GI biopsies and stool samples of infected individuals, pointing to a potential fecal-oral transmission route [10–12].

Animal models for COVID-19, involving non-human primates and other vertebrates, have reinforced the evidence of the virus’ ability to infect and replicate in the digestive system, leading to extended periods of viral shedding in feces [reviewed in [13]]. *in vitro* and *ex vivo* investigations, exploiting human intestinal epithelial cells, organoids, and human gut-on-chip models, have provided insights into SARS-CoV-2-induced GI pathogenesis [14–17]. These studies highlight critical aspects of the infection outcomes, such as intestinal cell damage, the elicitation of local immune and inflammatory responses, with pronounced alterations in gene expression profiles, emphasizing the need for comprehensive research to elucidate the dynamic processes responsible for the GI complications associated with SARS-CoV-2 infection.

In this framework, the fruit fly *Drosophila melanogaster* emerges as an invaluable model for exploring the pathogenesis of SARS-CoV-2. This is mainly attributed to the substantial genetic overlap between the fruit fly and humans, with around 90% of human proteins known to interact with the virus being conserved within the fly genome [18]. Recent advancements have led to the development of a *Drosophila* resource that enables tissue-specific co-expression of SARS-CoV-2 proteins alongside their corresponding human or fly binding partners, facilitating comprehensive *in vivo* analyses of viral-host interactions [19]. Earlier work demonstrated that transgenic flies expressing SARS-CoV-2 proteins such as Orf6, Nsp6, and Orf7, exhibit reduced survival rates and display significant anomalies in tracheal development and muscle architecture [18]. Particularity, ectopic Nsp6 expression causes increased glycolysis coupled to detrimental structural and functional alterations in the fly cardiac system, akin to some clinical manifestations observed in COVID-19 patients [20]. Furthermore, the application of Selinexor, a nuclear transport inhibitor, mitigates the adverse effects of viral protein Orf6, highlighting *Drosophila* potential for antiviral drug screening [18].

The current study leverages the *Drosophila melanogaster* midgut model to investigate SARS-CoV-2-induced intestinal pathophysiology. The *Drosophila* midgut shares structural and functional similarities with the human small intestine, serving as the primary site for food digestion and nutrient absorption [21]. Additionally, the intestinal epithelium acts as a barrier, safeguarding the organism from external invasive pathogens [22–24]. Our approach involved orally exposing flies to a clinical isolate of the original pandemic variant and characterizing its impact on the midgut homeostasis and physiology during the early stages of infection. Furthermore, our study demonstrated the effectiveness of this SARS-CoV-2 midgut model in evaluating the activity of promising anti-SARS-CoV-2 candidates such as Plitidepsin [25], and in establishing parallels with human pulmonary cells.

## Results

### SARS-CoV-2 administration by oral route compromises *Drosophila* survival

To assess the susceptibility of *Drosophila melanogaster* to SARS-CoV-2 *via* the oral route, wild-type mated females (*w^1118^*) aged 2-3 days, initially reared at 18°C, were shifted to 29°C for two additional days before virus ingestion and kept at this temperature during the experiments. This temperature was chosen to facilitate subsequent genetic manipulations and ensure the viability of the flies, while potentially maintaining the stability of the virus [26]. On the day of infection (D0), flies were starved for two hours to synchronize feeding behavior (Fig S1A).

The clinical isolate used for infection corresponds to the 2020 European D614G SARS-CoV-2 variant, derived from a nasopharyngeal swab and passaged once on Vero E6 cells (Fig 1A). The full-length genome sequence of this isolate is publicly available on GISAID under accession ID: RUN-PIMIT8 [27]. Infectivity of the viral stock was confirmed by immunostaining with an anti-SARS-CoV-2 spike-specific antibody (Fig S1B), and by monitoring cytopathic effects in Vero E6 cells (Fig S1C). For oral infection, flies were exposed to a dose of 0.45×10^6^ plaque forming unit (PFU) (Fig S1D), while mock-infected controls received cell culture supernatants.

**Fig 1.**
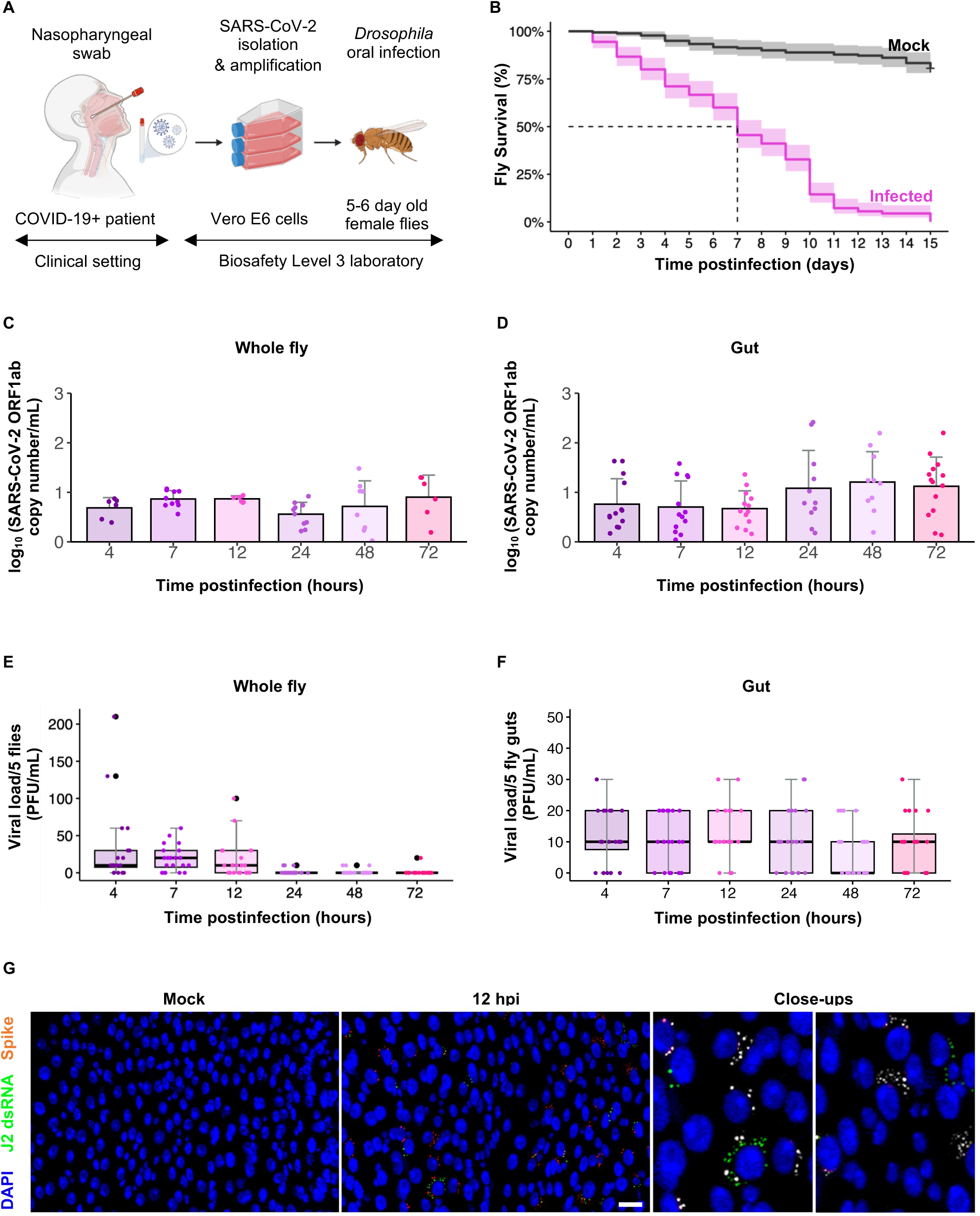
*Drosophila* susceptibility and permissiveness to SARS-CoV-2 oral infection. **(A)** Study design: 5–6-day old female flies were orally administered a solution of SARS-CoV-2, obtained from a nasopharyngeal swab of a COVID-19 patient. The clinical strain was isolated and amplified on Vero E6 cells. Created with BioRender. **(B)** Kaplan-Meier survival curves comparing flies (*w^1118^* genetic background) orally infected with SARS-CoV-2 to mock flies that ingested Vero E6 cell supernatants. Experiments represent three biological replicates, with 180 flies per condition in each replicate (total n= 540 flies per condition). Shaded areas represent the 95% confidence intervals (CI). Statistical significance was measured using the Log-rank test (p<0.01) for infected flies compared to mocks. **(C)** Longitudinal detection of SARS-CoV-2 genomic RNA using the Open Reading Frame 1ab (ORF1ab) RT-qPCR assay. Samples consisted of pools of 5 ground whole flies or guts **(D)**. Three independent experiments were conducted with 4 samples each (n=12 samples/condition/timepoint). Histograms show means with error bars indicating standard deviation. Statistical analysis using one-way ANOVA revealed no significant differences in viral RNA levels over time. (**E)** Longitudinal quantification of infectious viral loads using plaque forming assay (PFU). Samples consisted of pools of 5 ground whole flies or guts **(F)**. Three independent experiments were performed with 6 samples each (n=18 samples/condition/timepoint). Each dot represents values from one sample. (**G)** Representative images of a midgut confocal section dissected at 12 hpi, with its respective control. Midguts were stained with anti-SARS-CoV-2 spike antibody (red), anti-dsRNA J2 antibody (green), and DAPI (blue). Scale bar represents 50 μm. In the close-up views, Spike staining is shown in white to enhance contrast.

Our primary observations revealed a significant decline in survival rates of *Drosophila* orally infected with SARS-CoV-2 compared to the mock group, with a statistically significant difference in mortality (Log-rank test, p-value<0.001, Fig 1B). The risk of death for the infected flies increased over time, reaching a median mortality of 50% by day 7 postinfection and peaking to around 90% by day 10. Therefore, we focused mainly on the first 72 hours postinfection (hpi) in this study to capture the very early responses while fly survival is not highly affected, providing insights into the initial stages and progression of the disease.

To monitor viral presence, RT-qPCR was performed on total RNA samples from whole flies and dissected midguts. SARS-CoV-2 genomic RNA, targeting Open Reading Frame 1ab (ORF1ab) and the Nucleocapsid protein (N) genes, was detected as early as 4 hpi and persisted for at least 72 hpi (Fig 1C, 1D, S1E, S1F). Viral RNA levels showed no significant variation across the different time points (85% ± 10% of the samples testing positive; one-way ANOVA, p-value>0.5). Cycle threshold (Ct) average values reached 28 and 26 for ORF1ab and N, respectively. To assess the presence of infectious virus, PFU assays were conducted on homogenates from whole flies and isolated midguts. Despite technical challenges associated with applying infected fly homogenates on laboratory cultured cells, we successfully detected infectious particles in whole flies, with an estimated viral load of 15 PFU/mL during the early course of the infection (4-12 hpi) (Fig 1E). In midgut samples, we detected a mean viral load of 10 PFU/mL across all tested timepoints, in approximately 75% of the samples (Fig 1F).

Immunofluorescence staining for the Spike protein confirmed widespread distribution of viral antigen along the digestive tract, with a gut-wide prevalence of 72.2% ± 5.56% across the different timepoints (Fig 1G). To investigate replication activity, we employed the J2 antibody, which recognizes double-stranded RNA intermediates. Cells co-labeled with both Spike and J2 antibodies were observed, though not all signals colocalized (Figure 1G). Variability in nuclear morphology among labeled cells also suggests infection across multiple gut cell types.

These findings indicate that *Drosophila* exhibits susceptibility to SARS-CoV-2 following oral exposure, with measurable early mortality, persistent viral RNA, limited but detectable infectivity, and epithelial tropism. While the level of productive replication appears low, the consistent presence of viral RNA and virulence-associated mortality support the utility of *Drosophila* as a tractable *in vivo* model to study early-stage interactions between SARS-CoV-2 virulence factors and the GI epithelium.

### SARS-CoV-2 ingestion causes structural and physiological perturbations in the *Drosophila* intestine

Following oral exposure to SARS-CoV-2, pronounced morphological changes were evident in the midgut (Fig 2). Measurements of the midgut length, taken from the cardia center to the midgut/hindgut junction (Fig 2A and 2B), revealed a significant and constant decrease in size when compared to control flies, which exhibited a consistent midgut length of about 5.65 ± 0.15 mm. The reduction in midgut length began as early as 4 hpi and became more pronounced over time, eventually shrinking to 70% of its initial size (Fig 2B). Additionally, a significant increase in midgut width (Fig 2C) was observed starting at 7 hpi. Given the natural variability of midgut width along its length in normal conditions, width measurements were taken in a specific region in R4. These morphological alterations were coupled with intestinal obstructions, most notably at 48 hpi, where they were observed in 83% of the samples. These obstructions were characterized by the build-up of food materials in the hypertrophied crop (Fig 2D) and within the lumen of infected guts (Fig 2E), potentially interfering with normal food processing and impairing gut peristalsis. To further investigate gut motility, Phalloidin staining was used to visualize F-Actin filaments in the visceral muscles, revealing structural anomalies (Fig 2F). Notably, there was a significant increase in the thickness of the longitudinal syncytial muscle fibers throughout the midgut starting from 24 hpi (Fig 2F’), pointing to potential visceral muscle spasms. This thickening of muscle fibers, which persisted for at least 72 hpi (Fig 2F’), might be a contributing factor in the midgut shortening observed after SARS-CoV-2 infection. Alongside these structural modifications, SARS-CoV-2 ingestion led to a swift acidification of the entire digestive tract as early as 4 hpi (Fig 2G).

**Fig 2.**
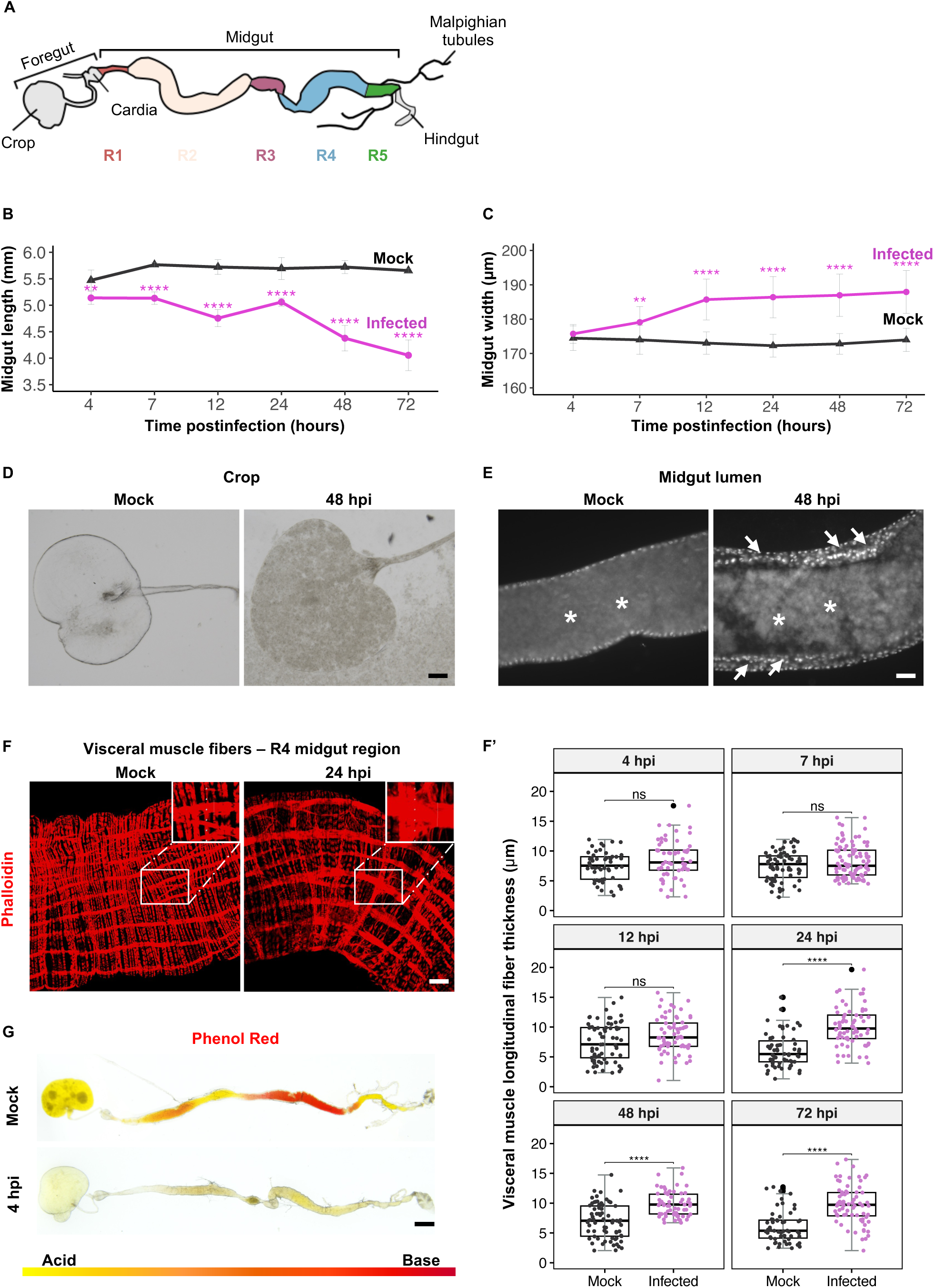
Effects of SARS-CoV-2 ingestion on midgut structure and pH in *Drosophila*. (**A)** Schematic representation of the adult *Drosophila* digestive tract. The midgut is subdivided into five functionally and morphologically distinct regions (R1-R5), each represented in a different color. (**B and C)** Measurements of gut length and width at different timepoints postinfection (n=18 midguts/condition/timepoint). Error bars indicate standard deviation. P-values correspond to the significant difference between conditions at the same time postinfection and were calculated using simple t-tests, corrected according to Bonferroni method. Significance levels are: ** P<0.01, **** P<0.0001. Absence of an asterisk indicates no significant difference (P≥0.05). **(D)** Representative images of crops from infected and mock flies at 48 hpi. Observations were performed across three independent experiments, with six crops analyzed per experiment (n = 18 crops per condition per time point). Scale bar represents 30 μm. **(E)** Images focusing on midgut lumens (R4 region; n=6, 3 replicates) of infected and mock flies. Guts were stained with DAPI at 48 hpi. Asterisks indicate the intestinal lumen, and arrows point to the thickened intestinal epithelium after infection. Scale bar represents 30 μm. **(F)** Longitudinal visceral muscle fibers (R4 region) of infected and mock flies, stained with Phalloidin (red) at 24 hpi. Scale bar represents 50 μm. (**F’)** Quantitative measurements of muscle fiber thickness across R4 region of dissected midguts at different times postinfection. Each dot represents measurements from a single longitudinal fiber. P-values are ns>0.05, **** <0.0001 calculated by Mann Whitney U test. **(G)** Full guts of wild type *w^1118^* flies fed with either mock or viral solutions mixed with Phenol red dye at 4 hpi (n=6 per condition, 3 independent replicates). Yellow indicates acidic regions (pH≤6), orange is neutral (6<pH<8), and red is alkaline regions (pH>8). Scale bar represents 350 μm.

Other organs, namely the rectum and ovaries, were also affected by SARS-CoV-2 infection (Fig S2). Approximately 90% of the examined rectums were hypertrophied, exhibiting a 200% ± 20% increase in surface area compared to mock-treated controls, along with a loss of clear segmentation in the rectal papillae, an alteration most prominent at 72 hpi (Fig S2A). Structural disorganization was also observed in the ovaries, which appeared reduced in size (Fig S2B). These findings further support the notion that SARS-CoV-2 infection in *Drosophila* extends beyond the gut and may impact multiple organs.

Together, these findings highlight the profound impact of SARS-CoV-2 enteric infection on the *Drosophila* model, characterized by notable structural and physiological changes within the digestive tract and other organs. These observations elucidate the complex nature of SARS-CoV-2 pathogenesis in this experimental system.

### SARS-CoV-2 leads to disruptions in the cellular composition of the *Drosophila* midgut

Having shown that SARS-CoV-2 impacts the structural integrity of the *Drosophila* alimentary canal, our investigation extended to the virus potential influence on the cellular makeup of the midgut epithelium, which appears thicker after infection (arrows in Fig 2E). This layer is primarily a single sheet composed predominantly of fully differentiated enterocytes (ECs) and enteroendocrine cells (EECs), which arise from progenitor cells known as enteroblasts (EBs) and pre-enteroendocrine cells (preEECs), respectively [28] (Fig 3A). To maintain tissue integrity and function under physiological conditions, multipotent intestinal stem cells (ISCs) sporadically divide, yielding another ISC and either an EB or a preEEC. These cells can be distinguished by their expression of specific transcription factors: ISCs, EBs, and preEECs express Escargot (Esg), EBs are marked by Suppressor of Hairless (Su(H)), and both preEECs and EECs express Prospero (Pros) (Fig 3A). In response to tissue damages, such as bacterial infection, this intestinal homeostatic program accelerates to effectively replenish lost cells [29]. The following section aims to discern whether SARS-CoV-2 infection disrupts this delicate cellular balance within the midgut epithelium.

**Fig 3.**
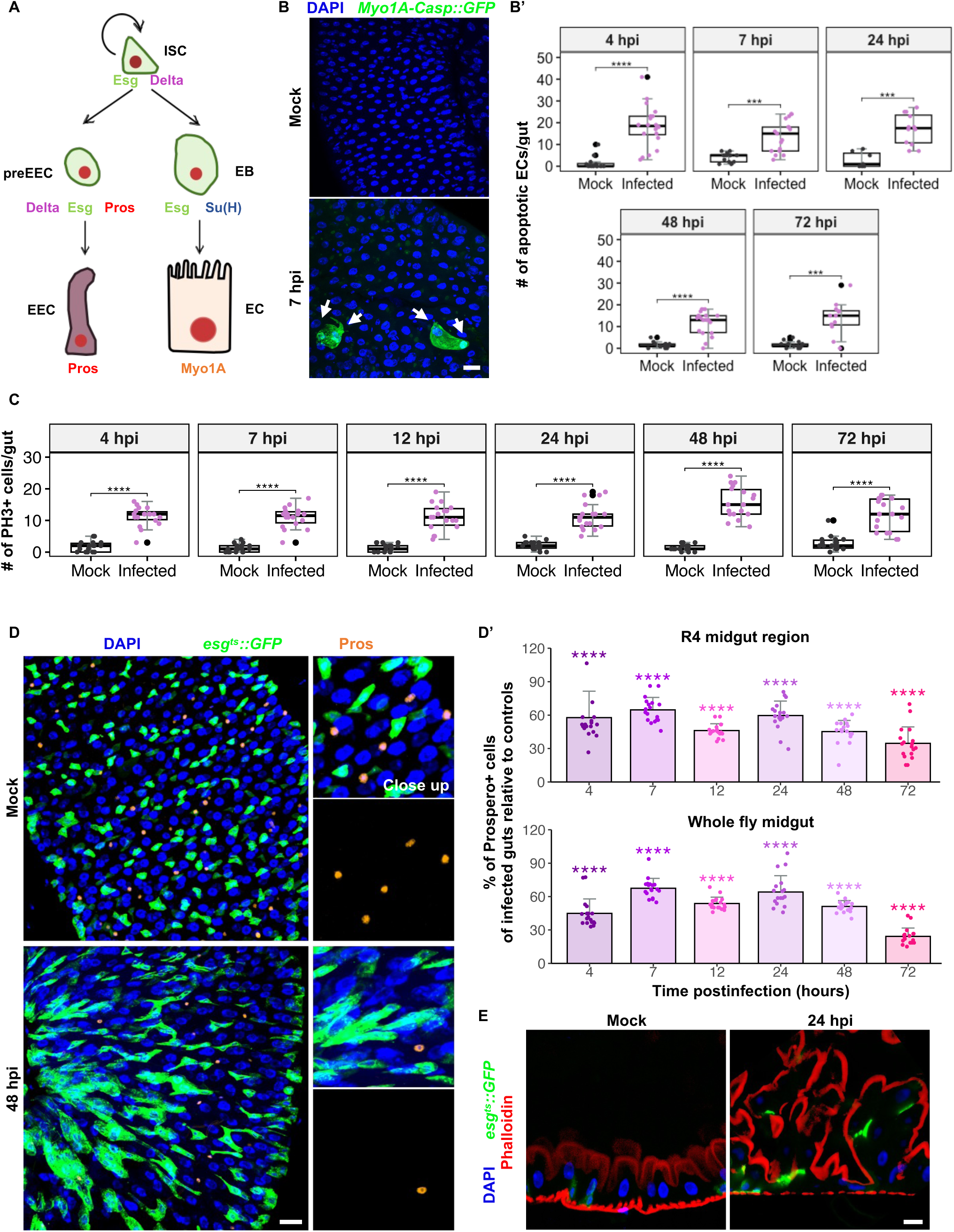
Disruptions in midgut cellular composition upon SARS-CoV-2 ingestion. **(A)** Schematic representation of the *Drosophila* adult intestinal stem cell lineage with cell specific markers. **(B)** Representative images of the R4 midgut region at 7 hpi showing Caspase 3 activity (GFP) driven by *Myo1A-GAL4* in ECs, compared to its respective mock. Arrows point to apoptotic ECs. Scale bar represents 50 μm. Quantification of apoptotic ECs per midgut at different times postinfection is shown in (**B’**). **(C)** Quantification of mitotic ISCs (PH3-positive cells) per midgut at different times postinfection. **(D)** Representative images of the R4 midgut region at 48 hpi and its respective mock, showing ISCs and EBs (*esg^ts^::GFP*, green), and preEE/EE cells (Pros, red). Scale bar represents 50 μm. Close-ups show merged and separate channels for Pros. (**D**’) Percentage of Prospero-positive cells in the R4 region and in whole infected guts relative to their controls. Histograms show means of the ratios, and error bars indicate standard deviation. **(E)** Confocal sections of *esg*-positive midguts expressing GFP in ISCs and EBs. F-actin in the brush border microvilli were stained using Phalloidin. Scale bar represents 30 μm. Data were collected from three independent replicates with 6 midguts each (n=18 midguts/condition/timepoint). In boxplots, each dot represents a count from one gut, large black dots mark outliers. P-values were calculated using simple t-tests corrected using the Bonferroni method. Statistical significance is indicated as *** P<0.001, **** P<0.0001. DAPI is blue.

SARS-CoV-2 is well-documented for its capacity to induce apoptosis in mammalian cell cultures. A recent investigation by Chu et al. [30] demonstrated that the virus strategically exploits caspase 6 in the apoptotic pathway to boost its replication efficiency. To assess the impact of SARS-CoV-2 on apoptosis specifically in ECs, the predominant midgut cell type, we used the *Myo1A>Casp::GFP* transgenic fly line. This line enables the expression of the Caspase 3 sensor, Casp::GFP, under the control of the *Myo1A-GAL4* EC-specific driver, as previously described [31,32]. In this system, GFP fluorescence is an indicator of Caspase 3 activation within ECs. Our analysis revealed an uptick in apoptotic ECs across the midgut regions of flies infected with SARS-CoV-2 compared to uninfected controls (Fig 3B). This increase in apoptotic activity was detected as early as 4 hpi and persisted at all subsequent timepoints (Fig 3B’).

Damage-induced cell death in the midgut usually triggers proliferation of ISC populations to maintain tissue integrity [33]. To test this hypothesis, we stained the midguts with an anti-PH3 antibody, which specifically marks phosphorylated histone H3 in condensed chromatin found in mitotic cells (S3A Fig). In the midguts infected with SARS-CoV-2, we spotted a 10-fold increase in mitotic indexes as early as 4 hpi, compared to their mock counterparts (Fig 3C). Interestingly, the proliferation rate remained consistently high up to 72 hpi. Concomitant with the elevated mitotic activity, there was a noticeable increase in the number of *esg^+^* progenitor cells (*esg^+^: esg*-*GAL4>UAS*-*GFP, tubP*-*GAL80^ts^*) throughout the midgut in response to SARS-CoV-2 enteric infection (Fig 3D). This augmentation was confirmed by quantifying the density of *esg^+^* cells within a defined 20,000 µm² area of the midgut R4 region, starting from 4 hpi to 72 hpi (Fig S3B). The high ISC activity in infected midguts was maintained in surviving flies at late days postinfection (Fig S3C and S3C’). Further confirmation of these findings came from observing an increase in the number of ISC-*Delta+* cells (*tubP GAL80^ts^; Delta GAL4>UAS GFP*) in SARS-CoV-2 infected midguts at 7 and 48 hpi compared to controls (Fig S3D and S3D’).

The GI tract functions as an endocrine organ, primarily through the activity of EECs that secrete hormones essential for various physiological processes including peristalsis, digestion, nutrient absorption, feeding behavior, and metabolism regulation [21]. Given this critical role, we investigated the impact of SARS-CoV-2 on EEC populations. Utilizing an anti-Prospero antibody to specifically mark preEE and EE cells, we found that SARS-CoV-2 enteric infection precipitates a significant decrease in the number of EECs in the midgut (Fig 3D). Notably, the EEC count was reduced by approximately 50% as early as 4 hpi and persisted until 72 hpi when compared to control midguts (Fig 3D’).

Taken together, these results demonstrate that SARS-CoV-2 enteric infection in *Drosophila* significantly disrupts intestinal cellular homeostasis, marked by increased apoptosis in ECs, elevated proliferation of ISCs, and a pronounced reduction in EEC numbers. This disruption is further evidenced by the altered structural organization and unusual multilayering of the intestinal epithelium (Fig 3E and schematized in S3E and S3E’ Fig), as revealed by Phalloidin staining that highlights the apical brush borders of ECs.

### SARS-CoV-2 impacts the replenishment of differentiated cells within the midgut epithelium

To thoroughly investigate the fate of ISCs after SARS-CoV-2 enteric infection, we performed cell lineage tracing experiments using the Repressible Dual Differential Stability Markers (ReDDM) genetic system [34]. The ReDDM methodology allows simultaneous expression of proteins with distinct half-lives: a short-lived mCD8::GFP and a long-lived H2B::RFP, under the regulation of cell specific drivers, including *delta-GAL4* (*Delta-ReDDM*), *Su(H)-GAL4* (*Su(H)-ReDDM*), and *esg-GAL4* (*esg-ReDDM*). This enabled us to monitor the dynamic cell behavior in the midgut over time as illustrated in Fig S4A, S4B, S4C.

Prior to infection, 3-day-old mated females were conditioned at 29°C for 3 days to activate cellular markers effectively. The *Su(H)-ReDDM* system highlighted a notable rise in the populations of *Su(H)^+^* (EBs: GFP^+^, RFP^+^) at both 4 and 48 hpi, compared to controls (Fig 4A and 4A’). Although the count of newly differentiated ECs (GFP^-^, RFP^+^) was higher than control levels at 4 hpi, this did not translate into a sustained EC replenishment at 48 hpi (Fig 4A’’). When using *Delta-ReDDM* genetic setup, *Delta^+^* cells (ISCs and preEECs: GFP^+^, RFP^+^) displayed an increase in number at 4 hpi (Fig 4B and 4B’), aligning with earlier findings using the *Delta GAL4^ts^>UAS GFP* fly line (Fig S3D). However, their progenies, encompassing EBs, ECs, and EECs did not show an increase in number at 4 hpi but demonstrated a significant rise in their numbers at 48 hpi (Fig 4B”), likely due to enhanced EB population as depicted with the *Su(H)-ReDDM* flies (Fig 4A and 4A’). Consistently, the *esg-ReDDM* system revealed a significant rise in the total count of *esg^+^* cells (ISCs and EBs: GFP^+^, RFP^+^) after infection compared to uninfected controls (Fig S4D and S4D’), yet without a simultaneous increase in the count of newly formed differentiated cells (GFP^-^, RFP^+^) (Fig S4D”).

**Fig 4.**
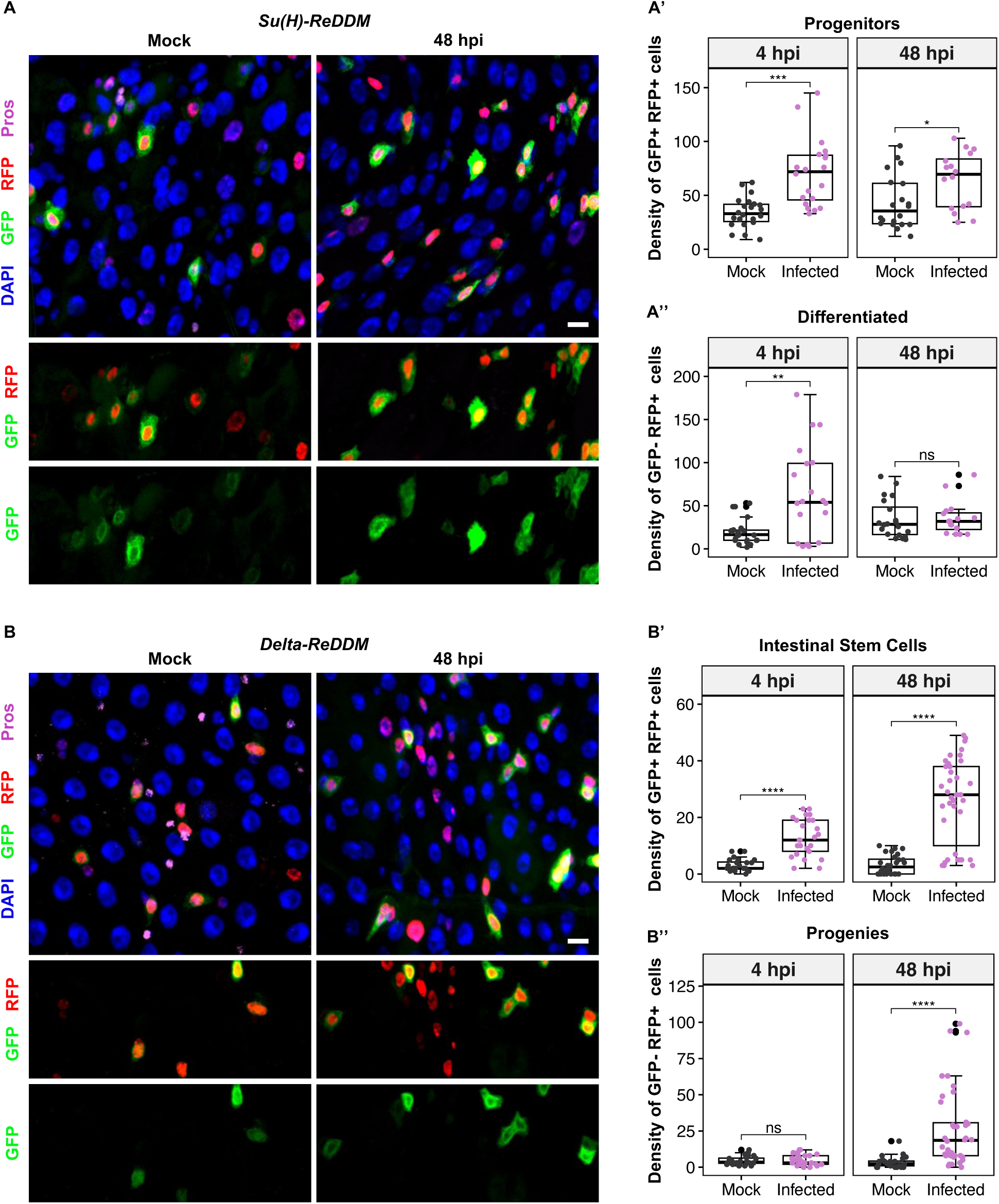
Impact of SARS-CoV-2 infection on midgut cellular regeneration and differentiation. **(A)** Representative images of *Su(H)*-*ReDDM* at 48 hpi. **(A’)** Quantification of the density of progenitors (EBs = Su(H)+ = GFP+ RFP+ cells) and (**A”**) differentiated cells (ECs + EEs = RFP+ only cells). **(B)** Representative images of *Delta*-*ReDDM* at 48 hpi. **(B’)** Quantification of the density of intestinal stem cells (ISCs (+ preEEs) = delta+ = GFP+ RFP+ cells), and **(B”)** their progenies (ECs + EEs + EBs = RFP+ only cells). DAPI is blue, and the scale bar represents 20 μm. Images were acquired at 48 hpi using confocal microscopy. Quantification was done in a specific surface area of 20,000 µm^2^ of R4 at 4- and 48 hpi. Data were collected from three independent replicates with 6 midguts each (n=18 midguts/condition/timepoint). Each dot represents count from one midgut and large black dots mark outliers. P-values from the Mann Whitney U-test are ns>0.05, * <0.05, ** <0.01, *** <0.001, **** <0.0001.

Overall, our data reveal that SARS-CoV-2 infection compromises the regenerative processes within the *Drosophila* midgut epithelium, affecting the replenishment of ECs and EECs, despite the induction of continuous ISC proliferation and increased EB numbers. The observed impediment in the transition to fully differentiated cellular states postinfection underscores the necessity for further investigations to unravel the mechanisms driving these disruptions.

### Transcriptomic profiling reveals a biphasic molecular response in SARS-CoV-2 infected midguts

To gain insights into the molecular dynamics of the response to SARS-CoV-2 enteric infection in *Drosophila*, we conducted comprehensive transcriptomic profiling at 4, 12, 24, and 48 hpi against mock conditions (Fig S5A and Table S1). Principal component analysis (PCA) indicated global shifts of midgut transcriptomes at all timepoints postinfection (Fig 5A). The first principal component (PC1) accounted for 48-59% of the total variance, while the second principal component (PC2) accounted for 24-33% of the variance. This separation along the PC1 and PC2 axes emphasizes the substantial global differences in midgut transcriptomes in response to SARS-CoV-2 infection over time.

**Fig 5.**
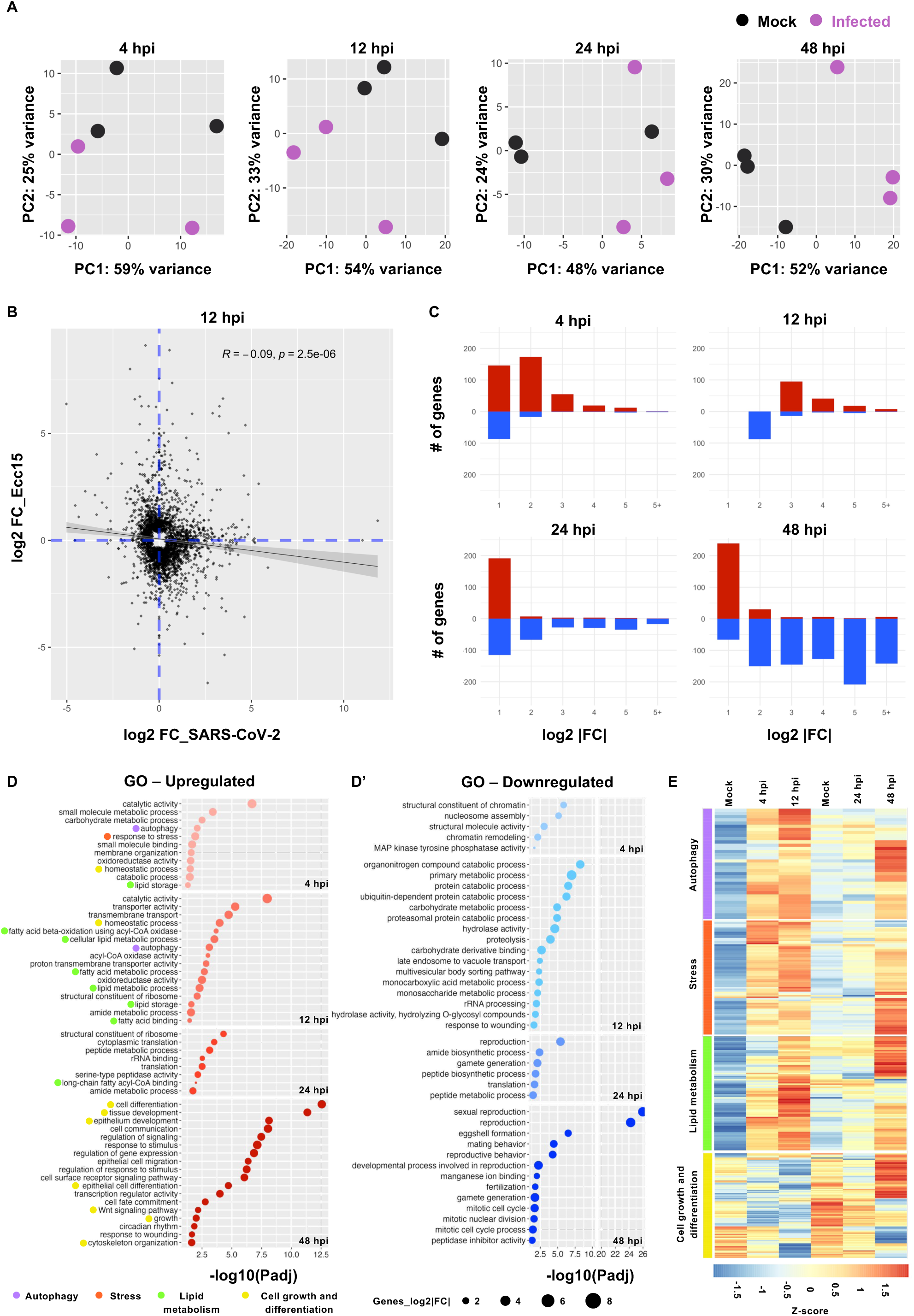
Biphasic molecular response to SARS-CoV-2 oral infection revealed by midgut transcriptomic profiling. **(A)** Principal Component Analysis (PCA) of the transcriptomes of SARS-CoV-2 infected guts vs. mock at different times postinfection. Three independent replicates were considered per condition. **(B)** Scatterplot comparing the log2 fold change of differentially expressed genes (DEGs) following *Erwina carotovora carotovora* (*Ecc15*) oral infection (y axis) and SARS-CoV-2 oral infection (x axis) at 12 hpi. **(C)** Transcriptomic variation in guts at different times postinfection, showing the number of upregulated (red) and downregulated (blue) genes categorized by the level of log2 fold change (log2FC). Only significantly DEGs (p-value<0.05) are shown in this graph. **(D)** Gene ontology (GO) enrichment bubble plot of upregulated and **(D’)** downregulated genes grouped by time postinfection. **(E)** Clustering heatmaps of gene expression across different times postinfection compared to their respective controls. Genes were clustered based on their GO categories.

Comparative analysis to midgut responses to pathogenic *Erwinia carotovora carotovora* (*Ecc15*) infection, which triggers robust induction of immune, stress, and developmental pathways [33], revealed distinct global expression profiles. Fold change values for genes at multiple timepoints (4, 12, and 48 hpi) following SARS-CoV-2 and *Ecc15* oral infection showed no correlation (*e.g.,* R^2^= −0.09, p= 2.5 e+06 at 12 hpi), indicating a specific transcriptomic response to SARS-CoV-2 (Fig 5B and S5B and Table S2).

Early response to SARS-CoV-2 infection (4 and 12 hpi) was predominantly characterized by upregulation of genes, with 2-4-fold increases being most common, and a relative scarcity of downregulated genes. As the infection progressed (24 and 48 hpi), the number of downregulated genes became more prominent, though some genes continued to show mild upregulation (less than 2-fold) (Fig 5C and S5C). A core set of genes consistently differentially expressed across timepoints emerged, with 307 genes upregulated and 60 downregulated at 4 and 12 hpi, versus 39 upregulated and 123 downregulated at 24 and 48 hpi (Fig S5D, S5D’, S5E and Table S3).

Gene Ontology (GO) analysis provided insights into the functions of differentially expressed genes (Table S4). Early upregulated genes (4 and 12 hpi) were predominantly associated with stress responses, autophagy, transmembrane transport, and lipid metabolism (Fig 5D). At later stages of infection (24 and 48 hpi), upregulated genes were mainly linked to translation (ribosomal constituents, rRNA binding), cell differentiation, epithelium development, growth, and cytoskeletal organization (Fig 5D). Downregulated genes at early timepoints were predominantly involved in chromatin remodeling, proteolysis, ubiquitin-dependent proteasomal protein degradation, and late endosome/multivesicular body transport (Fig 5D’). At later stages, downregulated genes included those involved in translation (a different set from the upregulated genes at earlier timepoints), reproduction, and mitotic cell cycle progression (Fig 5D’). We further examined the temporal changes in expression levels of specific gene categories critical during viral infection. Genes related to autophagy, stress responses, and lipid metabolism showed early upregulation, peaking at 12 hpi, followed by a decrease at 24 hpi and only a subset peaking again at 48 hpi (Fig 5E). In contrast, genes involved in cell growth and differentiation showed a decrease at early time points, peaking at high expression levels by 48 hpi (Fig 5E). The rapid upregulation of the highlighted host genes likely reflects immediate gut defense mechanisms while the virus is being internalized and replicating. The subsequent downregulation of key cellular processes at later stages may reflect the virus’s strategy to hijack the host’s cellular machinery, ensuring viral propagation.

In summary, our findings reveal a biphasic molecular response to SARS-CoV-2 enteric infection in the *Drosophila* midgut. Early responses are dominated by upregulation of genes related to stress response, autophagy, and lipid metabolism, whereas later stages exhibit downregulation of genes associated with translation and cell cycle progression. Understanding these molecular shifts provides crucial insights into the lifecycle of SARS-CoV-2 and its interaction with intestinal host biology, which may inform the development of effective antiviral strategies.

### SARS-CoV-2 induces region-specific disruptions in midgut lipid droplet organelles

Our transcriptomic analysis unveiled that SARS-CoV-2 promotes a marked transcriptional reprogramming of lipid metabolism genes in the early phase of enteric infection (Fig 5D and 5E). To better visualize the relationships between differentially expressed genes and the lipid metabolic pathways they are involved in, we performed a KEGG pathway enrichment analysis on the subset of genes modulated at 12 hpi, illustrated using a chordDiagram (Fig S5F). This representation highlights the overlap among genes contributing to multiple lipid-related processes, such as glycerophospholipid metabolism, fatty acid degradation, and sphingolipid metabolism. Notably, several genes were simultaneously implicated in distinct metabolic routes, suggesting coordinated regulation and potential functional convergence during infection. Building on this, we further explored the influence of SARS-CoV-2 on the homeostasis of intestinal lipid droplets (LDs), which are key cellular organelles involved in energy storage and lipid metabolism [35]. This investigation also relies on prior findings showing that SARS-CoV-2 induces LD accumulation in pneumocytes of COVID-19 patients [36]. Utilizing Nile Red and BODIPY stains, known for their efficacy in visualizing neutral lipids, we observed significant LD homeostasis disruptions within the *Drosophila* intestine at various postinfection intervals, notably at 7 and 12 hpi, as showcased in Fig 6A and S6A. These disruptions were heterogeneously distributed along the midgut, with pronounced, region-specific LD distribution variations (Fig 6A, 6B and S6A). Consistent with a previous study [37], our data confirm the natural abundance and size heterogeneity of LDs in the R2 region under uninfected conditions. After oral SARS-CoV-2 infection, however, there was a conspicuous depletion in both the number and size of LDs in this region, diverging markedly from the control. In the R3 and R4 regions, there was a noticeable accumulation of LDs postinfection. Intriguingly, even with the heightened LD density in the R4 region postinfection, most LDs were smaller in size compared to those typically found in R2 (Fig 6A). On the other hand, the R5 midgut region appeared to be unaffected by the virus-induced changes in LD profiles.

**Fig 6.**
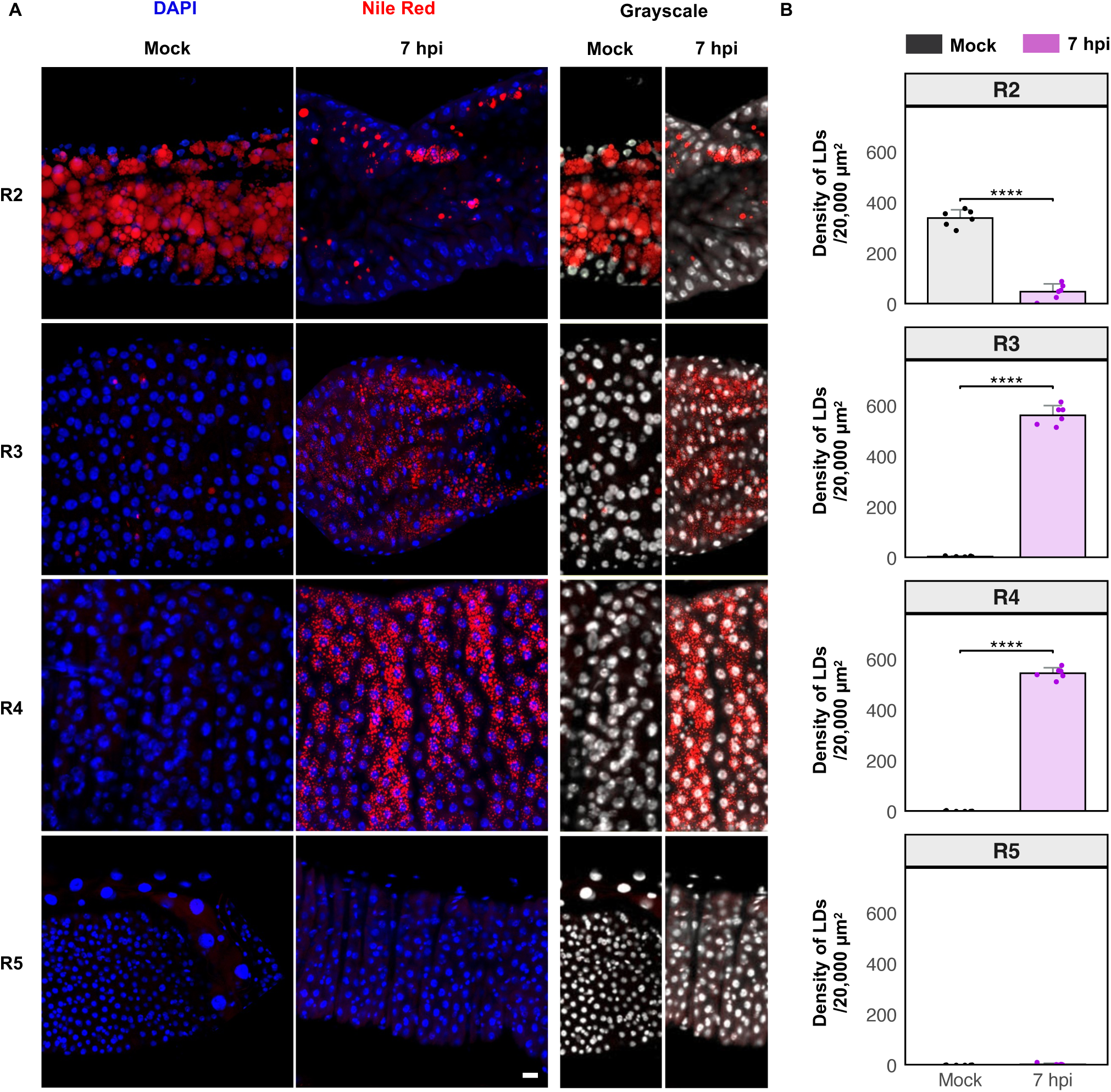
Region-specific disruptions in midgut lipid droplets homeostasis induced by SARS-CoV-2. **(A)** Representative images showing the distribution of lipid droplets in R2 to R5 regions of SARS-CoV-2 infected midguts at 7 hpi, using Nile Red staining. Comparisons are made to their respective mock samples. Nuclei are counterstained with DAPI (blue; white in grayscale panels) and scale bar represents 20 μm. A total of 20 midguts were scored for this experiment. **(B)** Density of lipid droplet particles in a 20,000 μm^2^ surface area of the R2-R5 midgut regions at 7hpi compared to mock midguts (n=6 per condition/region). Histograms show means and error bars indicate standard deviation. P-values were defined using individual t-tests and corrected using the Bonferroni method for multiple comparisons (ns P>0.05, and *** P<0.001).

These differential region-specific responses emphasize the complex interplay between SARS-CoV-2 and lipid metabolism which could impact the outcome of infection.

### Plitidepsin mitigates SARS-CoV-2-induced pathogenesis in the midgut

To explore potential antivirals effective against SARS-CoV-2-induced pathogenesis in the *Drosophila* midgut, we investigated the efficacy of Plitidepsin, a marine-derived cyclic depsipeptide isolated from the ascidian *Aplidium albicans*. Plitidepsin has shown potent preclinical efficacy against SARS-CoV-2 and has progressed to phase III clinical trials [25,38]. This drug was co-administered orally at different doses (0.01 µM, 0.1 µM, and 1 µM), both with and without the viral solution (Fig 7A). Initially, to establish the appropriate dose range of Plitidepsin for subsequent experiments, we evaluated its toxicity *in vivo*. Employing the *Myo1A>Casp::GFP* transgenic line, we observed a dose-dependent increase in apoptotic EC counts at 24 hours posttreatment (hpt), with a slight increase at 0.1 µM, which became excessive at 1 µM, indicating the high toxicity of this latter dose (Fig 7B and 7B’). We did not proceed to EC counts at 1 µM as the whole midgut becomes GFP positive. Similarly, we noted a significant dose-dependent increase in ISC proliferation, as evidenced by the rising number of PH3+ cells, quantified in Fig 7B”.

**Fig 7.**
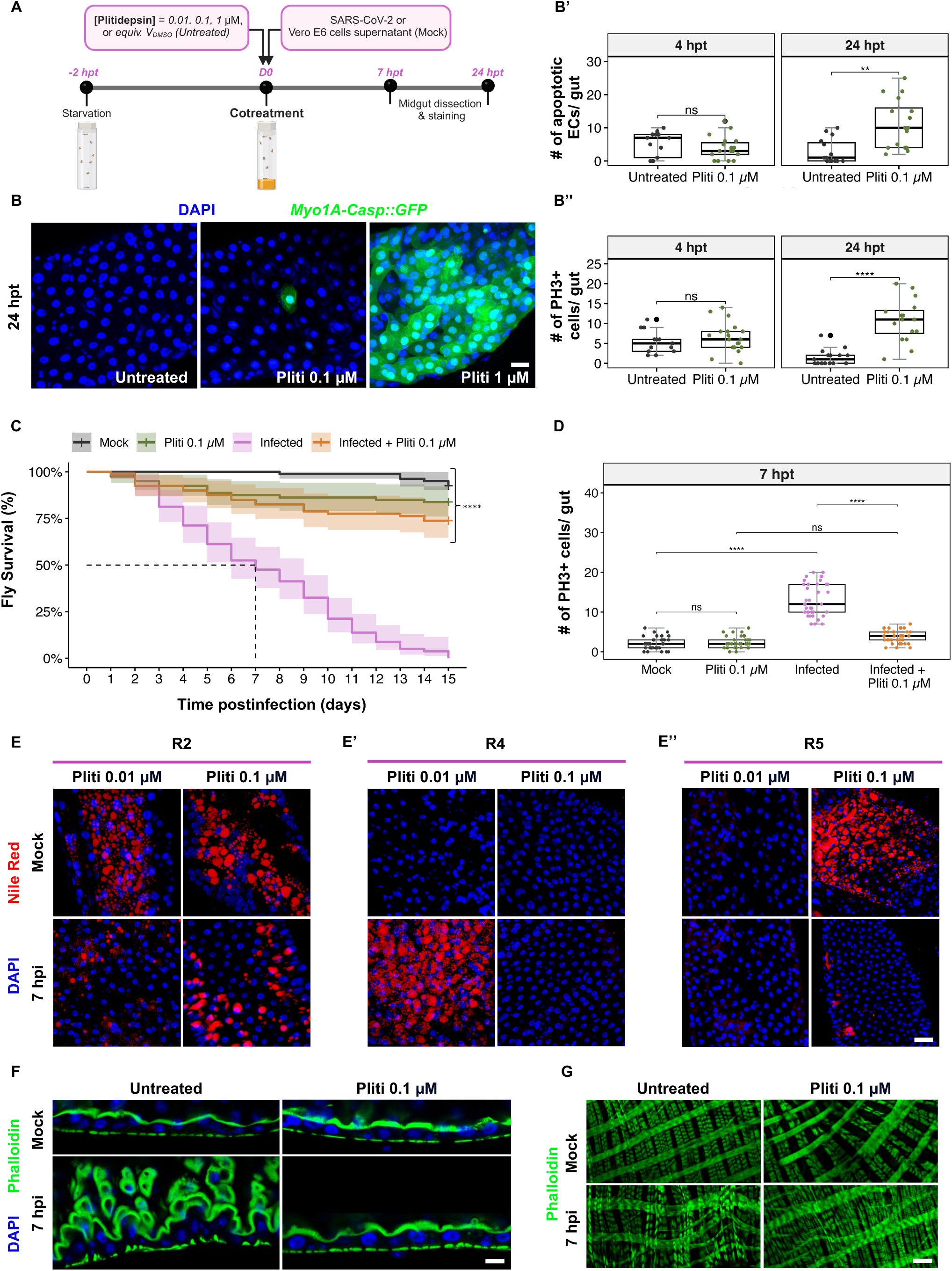
Plitidepsin mitigates SARS-CoV-2 activity in the midgut. **(A)** Diagram summarizing the cotreatment procedure of flies with different doses of Plitidepsin and SARS-CoV-2. Created with BioRender. **(B)** Representative images of R4 midgut region, showing Caspase 3 activity (GFP) driven by *Myo1A-GAL4* in ECs at 24 hours postingestion of 0.1 and 1 µM plitidepsin compared to untreated control. **(B’)** Quantification of apoptotic ECs (*Myo1A-Casp::GFP*), and **(B”)** mitotic ISCs (PH3-positive cells) in untreated guts or treated with 0.1 µM plitidepsin at 4 and 24 hours posttreatment. Data were collected from three independent replicates with 6 midguts each (n=18 midguts/condition/timepoint). P-values from one-way ANOVA are ns>0.05, ** <0.01, **** <0.0001. **(C)** Kaplan-Meier survival curves comparing flies (*w^1118^* genetic background) orally infected with SARS-CoV-2 or cotreated with 0.1 µM Plitidepsin, to mock flies that ingested Vero E6 cell supernatants or received 0.1 µM Plitidepsin alone. Data represent three biological replicates (n=20 flies/condition/replicate). Shaded areas represent the 95% confidence intervals (CI). Statistical significance was measured using the Log-rank test (p<0.01). **(D)** Quantification of mitotic ISCs (PH3-positive cells) in infected gut with SARS-CoV-2 or cotreated with 0.1 µM Plitidepsin, compared to mock flies that ingested Vero E6 cell supernatants or received 0.1 µM Plitidepsin alone. Data were collected from three independent replicates, with 12 midguts analyzed per condition in each replicate (n=36 midguts per condition). Statistical significance was assessed using one-way ANOVA. P-values: ns>0.05, **** <0.0001. **(E)** Representative images showing the distribution of lipid droplets at 7hpi in the anterior R2, **(E’)** in the posterior R4 and **(E”)** R5 regions, stained with Nile Red (red), following cotreatment with 0.01 µM and 0.1 µM Plitidepsin. The corresponding untreated control midguts are shown in Figure 6. 20 midguts were scored for this experiment. **(F)** Representative confocal sections of midguts showing the brush border microvilli at 7hpi following cotreatment 0.1 µM Plitidepsin, compared to untreated guts (n=10/condition/3 biological replicates). F-actin is stained using Phalloidin (green). DAPI is blue. Scale bar represents 30 μm. **(G)** Visceral muscle fibers in R4 at 7hpi, stained with Phalloidin (green) following cotreatment with 0, or 0.1 µM of Plitidepsin, compared to noninfected guts (n=10/condition/3 biological replicates).

Based on these results, the 1 µM dose was excluded from infection experiments due to its toxicity, and only non-toxic concentrations were used in subsequent analyses. Notably, treatment with 0.1 µM Plitidepsin significantly attenuated the SARS-CoV-2-induced lethality in flies (Fig 7C), supporting the efficacy of this dose in mitigating virus-induced pathogenesis.

We then assessed the effect of Plitidepsin cotreatment on specific intestinal alterations induced in the early phase (7 hpi) of SARS-CoV-2 infection, including ISC proliferation (Fig 7D), region-specific LD profiles (Fig 7E-E’’), epithelial cell multilayering (Fig 7F), and disturbances in visceral muscle fiber thickening (Fig 7G; as described earlier in Fig 6A and 3E and 2E respectively). At the cellular level, cotreatment with 0.1 µM Plitidepsin effectively counteracted the infection-induced rise in ISC proliferation (Fig 7D). At the metabolic level, 0.01 µM Plitidepsin had no observable impact on LD alterations caused by SARS-CoV-2 (Fig 7E-E” and S7A, controls in Fig 6); however, a 0.1 µM dose effectively restored LD distribution to that observed in uninfected controls in both R2 and R4 regions. Plitidepsin ingestion alone did not alter LD distribution in these midgut regions, regardless of the dose (Fig 7E and 7E’ and S7A and Fig 6). Interestingly, a 0.1 µM dose of Plitidepsin resulted in LD accumulation in the R5 region under non-infected conditions. We then investigated the potential of Plitidepsin in restoring the intricate architecture of the *Drosophila* intestinal epithelium caused by SARS-CoV-2 infection. At 0.01 µM, Plitidepsin began to counteract the virus-induced epithelial damage, partially restoring epithelial integrity. Importantly, 0.1 µM dose completely rescued the disrupted phenotype (Fig 7F). Nevertheless, Plitidepsin failed to exert any discernible rescue effect on the spasmodic status of visceral muscles at 7 hpi (Fig 7G). Intriguingly, Plitidepsin administration alone at 0.1 µM dose appeared to induce disrupted structuring of the visceral muscles all along the midgut compared to control conditions (Fig 7G).

These data provide supporting evidence for the therapeutic potential of Plitidepsin against SARS-CoV-2 *in vivo*, despite showing basal effect, and contributes to the growing body of research aimed at identifying effective therapeutics for combating COVID-19 and other viral outbreaks.

### Plitidepsin exhibits anti-SARS-CoV-2 activity and modulates lipid droplet accumulation in A549-ACE2 human pulmonary cells

Having demonstrated the modulatory effect of Plitidepsin on LD homeostasis in *Drosophila* SARS-CoV-2-infected midgut (Fig 7E and 7E’), we aimed to assess whether this metabo-modulating activity is conserved in a well-established human alveolar cell line that highly expresses ACE2 (A549-ACE2) for studying SARS-CoV-2 infection.

First, we confirmed that A549-ACE2 cells are permissive to the SARS-CoV-2 variant used in this study at a Multiplicity of Infection (MOI) of 0.1 (Fig S7B). Prior to assessing antiviral effects of Plitidepsin in A549-ACE2 cells, we examined its cytotoxicity using an ATP assay at 24- and 48 hpt, with concentration inhibiting 50% of cell viability (CC_50_) values of 0.88 and 0.02 µM, respectively (Fig S7C). This allowed us to evaluate the antiviral effect of Plitidepsin in these cells by flow cytometry, resulting in 50% inhibition concentration (IC_50_) of 0.003 µM at 24 hpi (Fig S7D). We further validated this antiviral effect by immunofluorescence assay (Fig S7E) using an anti-Spike antibody. Based on these results, we decided to carry out our subsequent experiments at an MOI of 0.1 and at Plitidepsin concentration of 0.01 and 0.1 µM, which are consistent with the doses used in the *Drosophila* experiments.

Strikingly, as observed in the R5 midgut region (Fig 7E”), our data indicated that Plitidepsin induces LD formation in non-infected A549-ACE2 cells in a dose-dependent manner, while reducing LD accumulation induced by SARS-CoV-2 at 24 hpi, as evidenced by Bodipy FLC12 LD-specific staining (Fig S7F and S7F’). This antiviral activity is coherent with our data obtained in the *Drosophila* midgut R2 and R4 regions (Fig 7E and 7E’).

Altogether, our findings in both the *Drosophila* midgut and human alveolar cells reveal a potential mechanistic link between Plitidepsin and LD homeostasis, highlighting the valuable use of the *Drosophila* midgut for unraveling region-specific metabolic modulation mediated by viral infection.

## Discussion

Although respiratory symptoms dominate the clinical presentation of COVID-19, increasing evidence highlights GI involvement, particularly in patients reporting digestive complaints. Imaging studies have identified COVID-19-associated gut abnormalities, including bowel wall thickening, pneumoperitoneum, and intestinal perforations [39]. Histopathological analyses of intestinal biopsies have further revealed epithelial alterations, such as cytoplasmic blebbing and cellular crowding in infected areas [40]. These observations underscore the need for tractable *in vivo* models to investigate the mechanisms of SARS-CoV-2-induced intestinal pathogenesis.

Leveraging the well-characterized *Drosophila melanogaster* midgut system [41], our study provides a multiscale *in vivo* dissection of the gut alterations triggered by SARS-CoV-2 infection. We demonstrate that oral exposure to SARS-CoV-2 can initiate a localized and functionally relevant infection in the fly midgut. Immunofluorescence staining using antibodies against the viral Spike protein and the replication intermediate marker J2 (dsRNA-specific) revealed intracellular viral signals within midgut epithelial cells. These signals were absent during the initial ingestion phase (1–3 hpi), supporting the interpretation that the detected Spike signal reflects active intracellular replication rather than passive retention of viral particles in the gut lumen. Despite the relatively low ingestion capacity of flies and the modest number of infectious particles recovered by plaque assay, viral RNA was consistently detected in midgut samples across multiple timepoints by RT-qPCR. This stable signal, coupled with immunostaining showing that up to 25% of epithelial cells display intracellular viral markers, suggests that viral replication, albeit possibly partial or abortive, is sufficient to disrupt epithelial homeostasis. Furthermore, the successful recovery of infectious virus from fly homogenates on Vero cells, despite technical limitations, supports the occurrence of productive replication in a subset of cells. Remarkably, this limited exposure was sufficient to induce rapid and severe intestinal remodeling. Within 24 hpi, flies exhibited structural abnormalities including gut shortening, epithelial and visceral muscle thickening, and luminal food retention. These phenotypes progressively worsened and ultimately culminated in fly death, likely driven by gut dysfunction and potentially exacerbated by systemic effects.

These early-onset changes seem to reflect an acute epithelial stress response involving both morphological remodeling and physiological disruption. Concomitantly, we observed an infection-induced shift in gut pH, which may result from direct viral effects or activation of innate immune pathways. At the cellular level, the emergence of a disorganized, multilayered epithelium, accumulation of apoptotic enterocytes, expansion of proliferative progenitors, and reduction of enteroendocrine cells all point to a rapid loss of epithelial homeostasis. This cellular destabilization appears to be part of a compensatory regenerative program. As early as 4 hpi, we observed a significant increase in EC apoptosis using the *Myo1A>Casp::GFP* reporter. Lineage-tracing with the *Su(H)-ReDDM* system revealed accelerated differentiation of ECs from the enteroblast pool, consistent with a rapid repair response to acute epithelial injury. In contrast, *esg-ReDDM* showed no net change in progeny numbers, likely reflecting a balance between enhanced EC production and concurrent EEC loss at this timepoint, confirmed by quantification of Prospero-positive cells. Although our study evaluated EC death, we cannot rule out EEC apoptosis. Attempts to assess EEC-specific death using *Prospero>Casp::GFP* proved inconclusive, likely due to the small size of EECs and difficulty distinguishing them from debris. We acknowledge this as a technical limitation and point to the need for other tools to dissect EEC turnover under viral challenge.

While the early response appears compensatory, our lineage-tracing data at 48 hpi reveal a more complex scenario. At this timepoint, all three systems, *esg-ReDDM*, *Su(H)-ReDDM* and *Delta-ReDDM*, show increased progenitor numbers, confirming sustained stem cell activation. Nevertheless, this expansion is not paralleled by efficient differentiation into mature intestinal cell types. In both *esg-ReDDM* and *Su(H)-ReDDM*, EBs appeared unable to fully commit to EC fates. However, the increase in *Delta-ReDDM* progeny likely reflects EB pool expansion, in line with *Su(H)-ReDDM* data. These observations suggest that SARS-CoV-2 infection sustains ISC proliferation while disrupting lineage differentiation, in addition to inducing cell death.

This decoupling between proliferation and differentiation is not typically observed in bacterial infections [42], which generally induce transient ISC activation followed by prompt epithelial restoration [43]. In contrast, our data support a model in which SARS-CoV-2 not only initiates epithelial turnover but also interferes with the proper execution of regenerative programs. This could reflect a general disruption of the differentiation machinery. Alternatively, it may represent a viral strategy to maintain a less immunocompetent, progenitor-rich environment. Importantly, such differentiation bottlenecks likely contribute to the epithelial disorganization and homeostatic collapse observed in infected midguts.

Although we did not directly address the molecular mechanisms underlying SARS-CoV-2 entry or identify the specific midgut cell types targeted by the virus, our findings lay important groundwork for addressing these questions. *Drosophila* harbors six conserved *ACE*-like genes [44], at least two of which, Ance and Acer, are expressed in the midgut (from flygut-seq database and [45]). These proteins have been proposed as potential orthologs of mammalian ACE2, the primary receptor for SARS-CoV-2. By integrating genetic tools and live imaging techniques available in *Drosophila*, further investigations could provide valuable insights into how SARS-CoV-2 interacts with the gut epithelial cells at the entry level.

In addition to structural and cellular remodeling, SARS-CoV-2 infection also induced early and significant midgut transcriptional reprogramming. Our transcriptomic profiling revealed that SARS-CoV-2 modulates the expression of a limited subset of genes in the midgut, suggesting a targeted rather than a broad response. The biphasic nature of the intestinal transcriptomic response is particularly noteworthy. The early activation phase is characterized by the predominance of upregulated genes and coincided with the onset of detectable viral replication, likely reflecting an immediate attempt by the host to counteract the infection. In contrast, the subsequent downregulation of transcriptional activity dampening key cellular processes, may denote an adaptive mechanism aiming to mitigate prolonged stress or defense responses. Interestingly, infected midguts revealed a pronounced upregulation of genes involved in lipid metabolism as early as 4 hpi, including those related to fatty acid β-oxidation, phospholipid remodeling, and lipid droplet dynamics. Gene ontology enrichment further highlighted deregulation in pathways such as glycerophospholipid metabolism and fatty acid elongation, suggesting that the enteric epithelium rapidly undergoes metabolic adaptation upon viral exposure. To assess the spatial and functional relevance of these transcriptional changes, we performed *in vivo* lipid staining using both Nile Red and BODIPY FL C12. Under mock conditions, LDs were largely restricted to anterior midgut regions. Upon infection, however, we observed striking reduction of LD in the R2 region and their accumulation in R4. These observations suggest that lipid metabolism is not only transcriptionally reprogrammed but also spatially reorganized, possibly as a result of viral manipulation or host compensatory responses. LDs are dynamic cellular storage organelles composed of neutral lipids, mainly triglycerides and cholesterol esters, enclosed within a phospholipid monolayer associated with various proteins [35]. While traditionally viewed as energy reservoirs, LDs also serve as hubs for lipid signaling and immune regulation and are increasingly recognized as key host platforms hijacked by intracellular pathogens. Viruses, in particular, have evolved strategies to exploit LDs as nutrient reservoirs or scaffolds for replication complexes [46]. Some lipid-enveloped viruses like coronaviruses and flaviviruses recruit LDs during their infection cycle by triggering their accumulation in host cells and modifying the lipid composition of these organelles to enhance viral replication [46]. Hence, LDs could serve as a platform for the virus to acquire lipids and components necessary for the assembly of new viral particles [47]. Viruses can also manipulate LDs for immune evasion by dissimulating within these particles to protect viral factors from recognition and degradation by the host immune systems [46]. Nevertheless, infected cells may also use LDs as part of their antiviral defense strategies. For example, some LD proteins, such as Viperin, have been implicated in viral restriction by interfering with viral assembly or triggering a variety of antiviral responses [48]. The differential regulation of LDs in the midgut could be linked to different metabolic functions or immune environments within the midgut. Functional interaction studies, along with profiling of lipid droplets and metabolic components across different gut regions, could provide insights into the metabolic requirements of SARS-CoV-2. Additionally, such studies may elucidate whether these antagonistic effects reflect a possible viral invasive strategy in the anterior midgut versus a lipid droplet-mediated cellular defense mechanism in the distal parts of the alimentary canal.

Comorbidities such as obesity and diabetes are known to exacerbate COVID-19 outcomes [49]. These conditions are often associated with altered lipid metabolism and chronic inflammation, which could further influence the viral pathogenesis observed in the gut. Through a screening of metabolic genes [50], a recent study found that key players like acetyl-CoA carboxylase-α (ACACA) and fatty acid synthase (FASN) can either inhibit or promote viral infection. Notably, the study highlighted the efficacy of Orlistat, an FDA-approved anti-obesity drug, as an inhibitor of FASN. Orlistat was shown to inhibit *in vitro* replication of SARS-CoV-2 variants and significantly reduced viral levels and lung pathology in a mouse model of SARS-CoV-2 infection, improving survival rates. Future research using available fly models of obesity and diabetes could help elucidate how these comorbidities influence the gut’s response to SARS-CoV-2 and whether targeting lipid metabolism, as demonstrated with Orlistat, might provide therapeutic benefits for patients with these conditions. This could pave the way for repurposing lipid metabolism inhibitors for the prevention and treatment of severe COVID-19, warranting clinical trials to evaluate their efficacy in humans.

Studies on other repurposed drugs like hydroxychloroquine and remdesivir revealed limited or no benefits in altering the disease progression [51]. Importantly, Plitidepsin, initially developed for multiple myeloma, has shown potent preclinical activity against SARS-CoV-2 [25]. This drug, which inhibits the eEF1A translation cofactor [25], shows promise in both animal studies and early human trials [38], though its full toxicity profile is pending. Interestingly, our results show that Plitidepsin mitigated SARS-CoV-2–induced gut phenotypes, suggesting antiviral activity in this model system. Despite its cytotoxic potential at higher concentrations, oral administration of non-toxic doses of Plitidepsin significantly improved fly survival in the context of SARS-CoV-2 infection. This phenotypic rescue highlights the utility of the fly model in assessing functional outcomes of viral exposure and therapeutic interventions.

Beyond its impact on mortality, Plitidepsin treatment attenuated several early infection-induced phenotypes in the midgut. Notably, it restored epithelial architecture, reduced the overactivation of intestinal stem cells, and normalized the distribution of lipid droplets, particularly in the R2 and R4 compartments where metabolic disruptions were most prominent. Although it does not restore the spasmodic visceral muscle phenotype, these effects suggest that Plitidepsin may limit viral replication and/or modulate host responses to restore homeostatic balance in infected tissues. Further investigation is necessary to determine whether Plitidepsin exerts its antiviral effects in the midgut through general translation inhibition or by regulating other cellular mechanisms, such as the modulation of lipid metabolism as observed in non-infected conditions both in *Drosophila* and human alveolar cells.

In conclusion, our study features the multifaceted impact of SARS-CoV-2 on the GI system, utilizing the *Drosophila melanogaster* model to reveal critical aspects of viral pathogenesis and host response. The findings emphasize the complexity of viral interactions with gut homeostasis, particularly in relation to tissue renewal and lipid metabolism. Additionally, our results demonstrate that *Drosophila* serves as a valuable model for *in vivo* screening of new bioactive compounds to combat COVID-19. Specifically, our study confirms the potential of Plitidepsin as a therapeutic agent against SARS-CoV-2, warranting further examination of its mechanism of action and therapeutic applications. Finally, given the continuous emergence of new SARS-CoV-2 variants, our research suggests that *Drosophila* can be invaluable in understanding how viral mutations impact intestinal pathogenesis. Together, our findings not only uncover new insights into SARS-CoV-2 gut pathogenesis but also establish a versatile model to accelerate antiviral discovery and dissect host-virus interactions at epithelial interfaces.

## Materials and methods

### SARS-CoV-2 clinical strain isolation

A SARS-CoV-2 strain was isolated from a nasopharyngeal swab taken from a tourist visiting La Reunion Island in April 2020. The individual had been confirmed positive for SARS-CoV-2 through RT-qPCR testing. The isolation process followed the methodology previously described by [27]. In brief, the African green monkey kidney cell line, Vero E6 (ATCC-CRL1586), was cultured at 37°C in a 5% CO_2_ atmosphere within a 25 cm² flask. The flask was filled with 5 ml of a solution containing Eagle’s Minimum Essential Medium (MEM, PAN Biotech, P04-08500) enriched with 5% heat-inactivated fetal bovine serum (FBS, PAN Biotech, P40-37500), 2 mmol/L L-glutamine, 1 mmol/L sodium pyruvate, 100 U/mL penicillin, 100 µg/mL streptomycin, and 0.5 µg/mL Amphotericin B (PAN Biotech, Aidenbach, Germany). One day later, the cells were washed with MEM containing 2% FBS, followed by the inoculation with 75 µl of the SARS-CoV-2 swab specimen. After a two-hour incubation, the inoculum was removed, cells were washed again, and replaced with fresh MEM 2% FBS. The flasks were then incubated and monitored daily for any cytopathic effects. Three days post-inoculation, supernatants were collected, centrifuged at 400 x g for 5 minutes at 4°C to remove cell debris, aliquoted, and then stored at −80°C.

The viral genome extracted from these samples was sequenced, with >90% coverage, using the MinION sequencing nanopore technology. The sequencing results confirmed the strain as belonging to the ancestral European D614G variant. Full genomic data have been made available on GISAID under the accession ID: RUN-PIMIT8. All experiments involving SARS-CoV-2 were conducted in a Biosafety Level 3 Laboratory (BSL-3) to ensure stringent containment and safety measures.

### SARS-CoV-2 production and titration

To produce a high-titer virus stock solution, 200 µL of the isolated SARS-CoV-2 was spread in permissive Vero E6 cells within 75 cm² flasks using MEM supplemented with 2% FBS. Cytopathic effect indicative of viral infection was observed three days post-inoculation, time at which viral lysates were collected. In parallel, supernatants from uninfected Vero E6 cells were harvested to serve as a mock solution. Following the protocol outlined previously, both types of supernatants were centrifuged, aliquoted, and stored at −80°C.

The concentration of viral progeny was determined using the PFU assay. For the assay, Vero E6 cells were plated in 24-well plates at a density of 70,000 cells per well. The next day, the cell monolayers were infected with 100 µL of a serial tenfold dilution of the SARS-CoV-2 solution, with the mock solution tested in parallel as a control. Two hours after allowing the virus entry, the cells were overlaid with 1% carboxymethyl cellulose (CMC, Sigma, C5013-500g) in MEM 5% FBS to restrict virus spread to directly adjacent cells. After incubating for three days at 37°C in a 5% CO_2_ atmosphere, the CMC overlay was removed. Cells were then washed twice with phosphate-buffered saline (PBS) and fixed with 3.7% paraformaldehyde (PFA, Thermo scientific, Cat# 043368.9M) in PBS for 20 minutes. Subsequent staining with 0.1% crystal violet (Sigma, C0775-100G) in 20% ethanol (Sigma, 24103-1L-R) allowed for visual identification of plaques as clear zones against a violet background of uninfected cells. After rinsing the plates with water to remove excess stain, plaques were counted and expressed as PFU/mL. All titrations were performed in duplicates. The viral titer was calculated using the formula: Titer (PFU/mL) = number of plaques counted x dilution factor/ volume of virus solution added to the wells (in mL).

To carry out PFU counts on flies, Vero E6 cells were infected with 100 μL of fly lysates diluted in MEM 2% FBS. Titration assays were conducted with at least five different replicates per experiment, and three independent biological replicates were performed. It was crucial not to fix the flies or midguts in paraformaldehyde prior to homogenization for this assay.

### Fly strains and crosses

The fly strains used in this study: *w^1118^* (BDSC, #5905), *w; esg-GAL4,UAS-GFP,tubP-GAL80^ts^* (*esg^ts^::GFP,* [52]), *w; tubP-GAL80^ts^*; *DL-GAL4,UAS-GFP/TM6b* (*Delta^ts^::GFP*, [53]), *w; Myo1A-GAL4/SM6b; UAS-GC3^AiG7s^/TM6b* (*Myo1A-Casp::GFP,* [31,32]), *w; UAS-GFP::CD8; UAS-H2B::RFP/TM2* (*ReDDM,* [34]), *w; Su(H)-GBE-GAL4, UAS-CD8::GFP/CyO; UAS-H2B::RFP, tubP-GAL80^ts^/TM6C* (*Su(H)-ReDDM,* [53]). Fly stocks were maintained at 25°C in polystyrene vials containing a standard culture medium (Bloomington recipe media, Nutri-fly, Cat#: 66-112) complemented with Amphotericin B (0.5 µg/mL). The flies experienced a regular 12:12-hour light-dark cycle and were transferred to fresh food vials weekly.

For lineage tracing studies, virgin females from the *Delta^ts^-GFP* and *esg^ts^-GFP* were crossbred with *ReDDM* males, whereas *Su(H)-ReDDM* fly stock were crossed with males possessing *w^1118^* genetic background. For all other experiments, virgin females were mated with *w^1118^*males. The different crosses were kept at 18°C. The resulting F1 progenies were housed in fresh medium for 2-3 days. Subsequently, mated females of the indicated genotypes were moved to 29°C for 2 days prior to infection.

### Fly infection and treatment

For oral viral infection, we subjected female flies to a 2-hour starvation period at 29°C in empty vials. These vials were prepared with standard culture medium completely covered at the top with a Whatman paper disk. Specifically, a 22 mm diameter disk was used, onto which was added 150 μL of either a Vero E6 cell supernatant (serving as the control, or "Mock" group) or a SARS-CoV-2 clinical isolate solution at 3.10^6^ PFU/mL. Following a 12-hour exposure period to these conditions, the flies were transferred to new standard food vials daily.

To evaluate the antiviral effectiveness of Plitidepsin (Pharmamar, PM90001) *in vivo*, starved flies were fed a solution containing either the virus or the mock solution, supplemented with different Plitidepsin concentrations (0.01, 0.1, and 1 µM).

### Fly survival assay

To evaluate the effects of oral SARS-CoV-2 infection and Plitidepsin treatment on *Drosophila* survival, flies were housed in groups of 20 per vial. For the infection assays presented in Figure 1, each experimental condition was represented by nine vials, and the experiment was independently replicated three times. For the Plitidepsin rescue experiments shown in Figure 7, each condition included three vials, with three independent biological replicates. Survival was monitored daily over a two-week period, and dead flies were removed and recorded throughout the experiment. To mitigate risks such as drowning in the food medium or fungal contamination, the vials were replaced daily.

### RNA extraction and RT-qPCR

For RNA extraction and subsequent RT-qPCR analysis, samples consisting of either five whole flies or five dissected midguts (excluding the Malpighian tubules) were homogenized in 500 µL of MEM supplemented with 2% FBS. Homogenization was achieved using a Tissue Lyser II, through 3 cycles of 1 minute each at a frequency of 30 Hz. The homogenates were then centrifuged at 4°C for 10 minutes at maximum speed. From each sample, 250 µL of the supernatant was utilized for RNA extraction, with the remaining volume being stored at −80°C for PFU assays. Total RNA was extracted using the RNeasy Mini Kit (Qiagen, Valencia, CA, USA) in accordance with the manufacturer’ instructions.

For cDNA synthesis, a two-step reverse transcription process was employed using ProtoScript II Reverse Transcriptase (Bioloabs) and Random Primer 6 (BioLabs). The reverse transcription conditions were as follows: initial denaturation at 70°C for 5 minutes, primer annealing at 25°C for 10 minutes, extension at 42°C for 50 minutes, and a final inactivation step at 65°C for 20 minutes. Quantitative real-time PCR (qPCR) targeting the Nucleoprotein (N) gene of SARS-CoV-2 was conducted using SYBR Green reagent (Thermo Fisher Scientific). Reaction mix included 12.5 μL of SYBR Green, 1 μL of each primer (1 μM), 5.5 μL of PCR water and 5 μL of cDNA. The qPCR assays were performed on a CFX96 Real-Time PCR Detection System (Bio-Rad), under the following conditions: activation at 95°C for 15 min and 40 cycles of denaturation and annealing/extension at 95°C for 15 s and 60°C for 1 min. Sequences of the primers used: Forward-HKUN 5’TAATCAGACAAGGAACTGATTA3’, Reverse-HKUN 5’CGAAGGTGTGACTTCCATG3’, Forward-ORF1ab 5’CTAGGACCTCTTTCTGCTCA3’, Reverse-ORF1ab 5’ACACTCTCCTAGCACCATCA3’. To establish a standard curve, cDNA synthesized from a SARS-CoV-2 positive sample previously verified in our laboratory was used as a positive control. Additionally, each sample was analyzed for the expression of the *Drosophila* housekeeping gene RpL32 to ensure accurate quantification. Sequences of RpL32 primers: Forward_RpL32 5’ATGCTAAGCTGTCGCACAAATG3’, Reverse_RpL32 5’GTTCGATCCGTAACCGATGT3’.

### Immunostaining

For midgut study, prior to tissue dissection in a BSL-3 containment facility, flies were initially anesthetized by placing them on ice for 1 minute. They were then gently rinsed in 70% ethanol for 2 minutes to ensure sterility and minimize contamination risk. The guts and ovaries were dissected out and fixed in 4% paraformaldehyde for 30 minutes at room temperature. Tissues were subsequently permeabilized with 0.1% Triton X-100 in PBS (PBST) for 5 minutes. Samples were blocked using 1% Bovine Serum Albumin (BSA) in PBST for 30 minutes at room temperature. Primary antibody incubation was carried out overnight at 4°C in a solution of PBST with 1% BSA. The primary antibodies utilized included: mouse anti-Prospero (DSHB Cat# MR1A, diluted 1:500), rabbit anti-Phospho-Histone H3 (Millipore Cat# 06-570, diluted 1:1000), mouse anti-dsRNA clone rJ2 (Merck Millipore, Cat# MABE1134) diluted 1:60), and human anti-Spike protein (Invivogen Cat# cov2rbdc1-mab1, diluted 1:1000). Following the primary antibody incubation, samples underwent three washes in PBST for 5 minutes each, then were incubated for 2 hours at room temperature with Alexa Fluor-conjugated secondary antibodies (Invitrogen, A11013, A21207, A11029, A-21235, A11014, diluted 1:1000). For cytoskeletal visualization, F-actin in both gut and ovary samples was stained using iFluor-conjugated Phalloidin (Phalloidin iFluor 488 Conjugate (1:1000), Abcam Ab176753 and Phalloidin iFluor 594 Conjugate (1:1000), Abcam Ab176757). Nuclei were stained with DAPI (Sigma, at 0.5 mg/mL). After staining, tissues were washed three times with

PBST and mounted on slides using a 1:1 mixture of PBS and glycerol. Gut samples were imaged using a NIKON C2si confocal microscope with NIS Elements software or Nikon Eclipse 80i fluorescence microscope, and ovary images were captured with a Nanozoomer S60 (Hamamatsu). All images were taken in the R4 midgut region, unless stated otherwise.

For *in cellulo* study, human alveolar cells overexpressing ACE2 (Invivogen, A549-ACE2) were regularly maintained in complete growth medium, supplemented with 0.5 μg/mL of Puromycin (Invivogen Ant-pr). Cells were plated on coverslips at a density of 100,000 cells/well. The following day, cells were subjected to either the mock, or the viral solution at different multiplicities of infection (MOI = 0.1, 0.2, 0.5, and 1). They were also cotreated with a ten-fold dilution series of Plitidepsin. 24 hours later, cells were rinsed with PBS, and fixed for 20 mins in PBS, PFA 3.7%. Next, cells were permeabilized with PBST for 5 mins, blocked with BSA 1% for 20 mins, and incubated then at room temperature with human anti-Spike protein (diluted 1:1000). Two hours later, cells were rinsed again with PBS and incubated for 30 additional minutes with Alexa Fluor 594 goat anti-Human IgG (diluted 1:1000). Nuclei were stained with DAPI. Two additional washes in PBS were performed and coverslips were mounted using 1:1 mixture of PBS and glycerol. Images were acquired using a NIKON C2si confocal microscope with NIS Elements software.

### Midgut morphometry

To assess midgut length and width, the guts were imaged using a Nikon SMZ18 stereoscope equipped with a Hamamatsu digital camera (Life Sciences, Japan). Length measurements were conducted by tracing a spline line along the midgut, from the center of the proventriculus to the junction between the midgut and hindgut, marked by the branching of the Malpighian tubules, whereas width was assessed by tracing a line throughout the R4 region. Measurements were taken for six guts per condition, and the process was repeated independently three times. For the observation of the crop and midgut lumen, a Nikon SMZ18 stereomicroscope equipped with a Hamamatsu digital camera was used (Life Sciences, Japan). Images were acquired from six samples per condition in each of the three experimental replicates. To measure muscle fiber thickness, the midgut was stained with Phalloidin, and the width of the longitudinal fibers was measured using ZenLite software. Measurements focused on sections from the posterior midgut in R4 region, with 10-13 fibers measured per gut across six guts for each condition.

### Midgut cell quantification

Mitotic cells marked for PH3 staining and apoptotic enterocytes were counted across all gut regions. Prospero+ cells were specifically counted in the R4 region. The enumeration was conducted using a Nikon Eclipse 80i fluorescence microscope. *esg^+^* cells (in *esg^ts^>GFP* line), *Delta^+^* cells (in *Delta^ts^>GFP* line), as well as both GFP^+^ RFP^+^ and RFP^+^ only cells (in *ReDDM* lines) were enumerated within a designated section of R4. Cell densities were calculated by normalizing the cell counts to a 20,000 µm^2^ surface midgut area using Zen software (ZEN Digital Imaging for Light Microscopy). Data were collected from six midguts per condition for each replicate, and the independent experiments were conducted a minimum of three times to ensure reliability.

### Lipid droplet staining

Guts dissected for lipid droplet staining were first fixed in 4% paraformaldehyde in PBS for 30 minutes, followed by 3 rinses in PBS. They were then permeabilized with 0.1% PBS-Triton X-100 for 5 minutes. For staining, the guts were incubated with a freshly prepared 1 µM solution of Nile Red (Sigma, Cat#: 72485) in PBST for 5 minutes or with an Oil Red O (Sigma, O0625-25g) working solution for 20 minutes. The Oil Red O working solution was prepared fresh as a 6:4 dilution of a 0.1% stock solution in isopropanol with distilled water, then filtered through a 0.45 µm syringe filter. After staining, the guts were washed twice and mounted in a PBS: Glycerol (1:1) mixture. Midguts and A549-ACE2 cells were stained with Bodipy FL C12 according to manufacturer’s instructions (Invitrogen, L3483). Imaging for Nile Red and Bodipy FL C12 staining was performed using a Nikon C2si confocal microscope with NIS Elements software. Mean fluorescence intensity of both dyes was quantified using ImageJ 1.52a, from image stacks containing the same number of slices, and normalized to DAPI signal.

### Midgut pH assay

To assess changes in gut acidity following SARS-CoV-2 infection, flies were fed a solution containing 0.1% (w/v) Phenol Red (Sigma), mixed with either mock or viral suspensions. Flies were starved prior to feeding to enhance ingestion efficiency. Successful dye ingestion was confirmed by the visible presence of red coloration in the abdomen within minutes of exposure. Only flies with clear abdominal dye accumulation were selected for subsequent analysis. At 4 hpi, flies were dissected, and midguts were immediately imaged using a Nikon SMZ18 stereomicroscope. As gut pH is highly sensitive to dissection and rapidly changes *ex vivo*, all imaging was performed within minutes to preserve physiological pH profiles.

### Gut RNA-seq analysis

Following the oral virus infection steps, 12 guts per replicate including the crop, midgut, hindgut, and rectum were dissected in sterile RNAse free PBS. The study examined multiple timepoints: 4, 12, 24, and 48 hpi, with three independent replicates for each condition.

Like viral infection protocol, bacterial oral infection was carried out at 29°C by using vials in which the fly food was covered by a Whatman filter paper containing 150 µl of either 2.5% sucrose solution (control), or mixed solution of 75 µl 5% sucrose with equal volume OD600 = 200 bacterial pellets. Midguts were dissected in PBS 1X at 4, 12, and 48 hpi.

Samples were homogenized in RNAse-free tubes with 500 µL of TRIzol and subsequently stored at −80°C. Next, total RNA was extracted via a modified phenol-chloroform method [54]. Briefly, samples were thawed on ice and 120 µL of chloroform were added. Samples were vortexed and centrifuged for 15 min at 12,000X g, at 4°C. Next, the aqueous supernatant phase was transferred to a fresh tube, where it was mixed with 700 µL Buffer RLT (Qiagen RNeasy kit, cat #74004), and then with 500 µL of 100% ethanol. 700 µL at a time of the mixture were passed through a RNeasy spin column and centrifuged for 20 s at 10,000X g, followed by a final spin for 1 minute at 10,000X g to dry the column. The membrane was then washed twice with 500 µl of RPE and centrifuged for 20 s at 10,000X g, and centrifuged for 1 minute at 16,000X g to dry the column. Samples were eluted in 30 µl of RNase-free water.

Libraries were prepared using the Quantseq 3’ mRNA-seq prep kit from Lexogen according to the manufacturer’s instructions. Sample quality was evaluated before and after library preparation using a fragment analyzer (Advanced Analytical). Libraries were pooled and sequenced on the Illumina Nextseq 500 instrument using standard protocols for 150 bp single end read sequencing. Sequencing was carried out at Cornell University BRC Genomics Core Facility (RRID:SCR_021727). Raw sequence data have been deposited at Sequence Read Archive (SRA PRJNA1137395 for *Ecc15* and SRA PRJNA1137405 for SARS-CoV-2) and are publicly available. Reads were mapped to the host transcriptome (r6.55) using Salmon and differences in expression levels after infection were calculated using DESeq2 in R. Gene Ontology over-representation analysis was performed using g:Profiler [55]. Kyoto Encyclopedia of Genes and Genomes (KEGG) pathway enrichment analysis was performed on the subset of significantly modulated genes involved in lipid metabolism related GOs using DAVID online ressources. PCA plots, Volcano plots, Venn diagrams, GO bubble plots, correlation plots, clustering (heatmap) of gene expression, and chordDiagrams were generated in R using ggplot2, DEseq2, pheatmap, and circlize packages. Transcriptomic profiles from 4 hpi and 12 hpi samples were compared to 4 hpi mock controls, while data from 24 hpi and 48 hpi samples were compared to 24 hpi mock controls.

### Cytotoxicity assay

Cell viability following Plitidepsin treatment was assessed *in vitro* using ATP assay (CellTiter-Glo 2.0 Cell Viability Assay from Promega). This assay was performed on A549-ACE2 cells. Cells were cultured in 96-well plates at a density of 2×10^4^ cells per well in DMEM 10% FBS. The following day, ten-fold dilutions series of Plitidepsin ranging from 10 µM to 10^-6^ µM were used to treat cells. At 24- and 48-hours posttreatment, cells were rinsed with PBS, and 100 µL of culture medium mixed with 50 µl ATP reagent. After 2 min incubation time, cells were mechanically lysed, and ATP release was revealed by measuring luminescence.

### Flow cytometry assay

Cells were rinsed twice with PBS 1X, detached with 20 µL of trypsin/EDTA, and then fixed with 3.7% paraformaldehyde in MEM 5% FBS solution for 15 minutes. After PBS wash, the cells were next incubated for 1 hour with the monoclonal human IgG1 anti-Spike antibody (Invivogen), diluted in 0.1% PBS-Triton X-100. Detection of the primary antibody was accomplished using a secondary goat anti-Human IgG antibody conjugated with Alexa Fluor 488 (Thermo Fisher Scientific), allowing the identification of SARS-CoV-2-infected cells. Following washing with PBS-Triton X-100, samples were subjected to flow cytometric analysis using Cytoflex (Beckman Coulter, Brea, CA, USA). Quantitative analyses were realized using CytExpert software.

### Statistical analysis

Data analysis and graph generation were performed using RStudio (R Statistical Software v4.2.0). At least, three biological replicates were performed for all experiments. Quantification data were first assessed for normality (Shapiro test) and homogeneity (Bartlett’s test). Results were then analyzed using one-way or two-way ANOVA followed by Tukey’s posthoc-tests, or Kruskal-Wallis followed by Dunn’s posthoc-test, Mann-Whitney U-test, or Student’s t-test (ns p>0.05, *p < 0.05, **p < 0.01, ***p < 0.001, ****p < 0.0001). Boxplots include 0.25, 0.5, 0.75 quantiles and mean represented by black lines. Line graphs and histograms were expressed as mean ± SD. The 50% cytotoxic (CC_50_) and inhibitory (IC_50_) concentrations were estimated using nonlinear regression to generate sigmoidal dose-response curves. For IC_50_, values are normalized against control groups of infected but untreated cells.

## Supporting information

**Table S1.** List of significantly differentially expressed genes upon SARS-CoV-2 oral infection.

**Table S2.** Correlation analysis of genes regulated following SARS-CoV-2 versus *Ecc15* oral infection.

**Table S3.** List of core upregulated and downregulated genes upon SARS-CoV-2 ingestion.

**Table S4.** Gene Ontology enrichment analysis of genes modulated by SARS-CoV-2 in the midgut.

## Acknowledgements

We thank Armel Gallet, Marie-Lise Gougeon, Philippe Despres, and Radwan Kassir for their valuable insights on this research. We are grateful to Cyrille Lebon and Hervé Pascalis for ensuring optimal experimental conditions at the PLATIN-OI BSL3 platform. Library sequencing was carried out at Cornell University BRC Genomics Core Facility (RRID:SCR_021727). This study was funded by POE FEDER 2014-20 of the Conseil Régional de La Réunion (TFORCE-COVIR, N°20201437-0027601), and Research Federation BioST, Université de La Réunion (COVIPID project). LEK received doctoral fellowships from the National Council for Scientific Research of Lebanon (CNRS-L), the Lebanese University and ERASMUS+MIC program. Buchon’s laboratory was supported by NIH R01AI148529, R01AI148541 and NSF IOS2024252. Library sequencing was carried out at Cornell University BRC Genomics Core Facility (RRID:SCR_021727).

## Author Contributions

Project Setup, L.EK., P.M., C.EK., and D.O.; Methodology and Validation, L.EK., C.EK., and D.O.; Investigation, L.EK., J.G., C.EK., and D.O.; RNA-seq library preparation and Raw Data Curation: PN and NB; RNA-seq Formal Analysis: L.EK., P.N., N.B., C.EK., and D.O.; Writing - original draft: L.EK. and D.O.; Writing – Review & editing, all authors. Supervision and Funding Acquisition, P.M., C.EK., and D.O. Conceptualization and Project Administration, C.EK. and D.O.

## Supplemental figure legends

**Fig S1. *Drosophila* oral infection approach with an infectious SARS-CoV-2 clinical isolate**

(A) Diagram summarizing the global approach to SARS-CoV-2 enteric infection in *Drosophila*. Oral infection occurs on day 0 (D0), with flies feeding on viral solution for 12 hpi before being transferred to normal medium. Fly preparation starts 5 days prior to infection (−5 dpi), and some experiments last till 14 days postinfection (14 dpi). Created with BioRender.

(B) Immunohistochemical staining of Vero E6 cell monolayers with anti-SARS-CoV-2 spike antibody (green) and DAPI (blue). Cells were observed under 10x magnification using fluorescent microscopy at 48 hpi. The top row shows mock or uninfected cells, while the bottom row shows cells infected with a multiplicity of infection (MOI) of 0.1. Scale bar represents 10 μm.

(C) Cytopathic effect observed in Vero E6 cells infected with a SARS-CoV-2 clinical isolate. Cell cultures were observed under 10x magnification using light microscopy at 72 hpi. The top row shows uninfected cells, while the bottom row shows cells infected with SARS-CoV-2 at an MOI of 0.1. Scale bar represents 10 μm.

(D) Quantification of SARS-CoV-2 viral production by PFU assay. Images display plaque morphology, size, and halo structure, characteristic of SARS-CoV-2. Two representative images are presented: the top row shows the 10^-4^ dilution, and the bottom row shows 10^-5^ dilution of the viral stock.

(E) Longitudinal detection of SARS-CoV-2 genomic RNA using the HKU Nucleocapsid (HKUN) RT-qPCR assay. Samples consisted of pools of 5 ground whole flies or guts **(F)**. Three independent experiments were conducted with 4 samples each (n=12 samples/condition/timepoint). Histograms show means with error bars indicating standard deviation. Statistical analysis using one-way ANOVA revealed no significant differences in viral RNA levels over time.

**Fig S2. SARS-CoV-2 induces structural disruptions in multiple *Drosophila* organs**

(A) Rectums of infected flies observed under a stereoscope showed structural aberrations at 72 hpi compared to control. The dotted lines in the infected rectum delimit the rectal papillae. Scale bar represents 20 μm.

(B) Ovaries of infected flies observed under a Nanozoomer microscope at 72 hpi showed a disorganized and atrophic structure compared to control. Stained with DAPI (blue) and phalloidin (red). Scale bar represents 100 μm.

**Fig S3. Expansion of intestinal stem cells after SARS-CoV-2 enteric infection in *Drosophila***

**(A)** Representative images of the R4 midgut region at 48 hpi, showing ISCs and EBs (*esg^ts^::GFP*, green), and mitotic figures (PH3, red). Arrows point to PH3-positive cells.

**(B)** Density of esg-positive cells in a 20,000 μm^2^ surface area of the R4 midgut region of SARS-CoV-2 infected flies and their controls at different times postinfection.

**(C)** Representative images of the R4 midgut region at 7 days postinfection (dpi), showing ISCs and EBs (*esg^ts^::GFP*, green), and mitotic figures (PH3, red). Arrows point to PH3-positive cells. (**C’**) Quantification of mitotic ISCs (PH3-positive cells) per midgut at 7 and 14 days postinfection in surviving flies.

**(D)** Representative images of the R4 midgut region at 48 hpi and its respective mock, showing ISCs (*Delta^ts^::GFP*, green) and mitotic figures (PH3, red). White arrows point to PH3-positive cells. (**D’**) Quantification of *Delta*-positive cells in the R4 region of SARS-CoV-2 infected flies and their respective controls at 7 and 48 hpi.

**(E)** Schematic diagram representing the structure of the intestinal epithelial monolayer and the major morphological changes observed after SARS-CoV-2 infection (**E’**). The basal membrane (BM) is represented in black, and the visceral muscle (VM) in purple. Dying cells are represented by dotted of segmented lines.

Data were collected from three independent replicates with 6 guts each (Total n=18 midguts/condition/timepoint). Each dot represents count from one gut. Large black dots mark outliers. P-values from simple t-tests corrected upon Bonferroni method (B, C’) are **** <0.0001.

**Fig S4. Lineage tracing systems for monitoring intestinal cellular turnover following SARS-CoV-2 infection**

Schematic diagram of the intestinal lineage tracing systems used in this study:

(A) *Su(H)*-*ReDDM*,

(B) *Delta*-*ReDDM*,

(C) *esg*-*ReDDM*.

The ReDDM (repressible dual differential stability markers) relies on the differential stabilities of a pair of fluorescent proteins: the short-lived membrane tethered mCD8-GFP (green), and the long-lived histone tethered H2B::RFP (red).

(D) Representative images of *esg*-*ReDDM* at 48 hpi. Quantification of progenitors (D’) (ISCs + EBs + preEEs = esg+ = GFP+ RFP+ cells) and (D”) differentiated cells (EBs + ECs + EEs = RFP+ only cells).

DAPI is blue, and the scale bar represents 20 μm. Images were acquired at 48 hpi under 40x magnification using confocal microscopy. Quantification was done in R4 at 4- and 48 hpi. Data were collected from three independent replicates with 6 midguts each (n=18 midguts/condition/timepoint). Each dot represents count from one midgut; large black dots mark outliers. P-values from the Mann Whitney U-test are ns>0.05, * <0.05, ** <0.01, *** <0.001, **** <0.0001.

**Fig S5. SARS-CoV-2 induces dynamic changes in *Drosophila* midgut transcriptome**

(A) Schematic diagram of the experimental workflow of the comparative transcriptome analysis.

(B) Scatterplot comparing the log2 fold change of differentially expressed genes (DEGs) between *Erwina carotovora carotovora* (*Ecc15* - y axis) and SARS-CoV-2 (x axis) at 4 and 48 hpi.

(C) Volcano scatter plot of differentially expressed genes at different times postinfection. Black dots represent genes where −1<log2 FC<1, red dots are significantly overexpressed genes (log FC>1, P-value<0.05), blue dots are significantly downregulated genes (log FC<-1, P-value<0.05), and gray dots are genes that are non-significantly different (P-value>0.05).

(D) Venn diagrams depicting the number of upregulated or (D’) downregulated genes showing significant differential regulation (p-value<0.05) at different times postinfection.

(E) Table showing the number of genes in the intersection that are significantly differentially expressed at different times postinfection. Pink-shaded cells highlight consistently changing genes.

(F) Chord diagram showing the association between differentially expressed genes and lipid metabolic pathways during early SARS-CoV-2 enteric infection. The diagram illustrates the connections between individual genes (left) and their associated metabolic pathways (right). Colors represent distinct lipid-related processes.

**Fig S6. SARS-CoV-2 modulates lipid droplet distribution in *Drosophila* midgut**

(A) Representative confocal images of *w^1118^ Drosophila* midguts stained with BODIPY^TM^ FL C12 dye at 12 hpi compared to mock. Scale bar represents 20 μm.

**Fig S7. Plitidepsin counteracts SARS-CoV-2 activity in *Drosophila* midgut and human A549-ACE2 pulmonary cells**

(A) Density of lipid droplet particles in a 20,000 μm^2^ surface area of the R2-R5 midgut regions (related to Figure 7E-E”) at 7hpi following cotreatment with 0.01 and 0.1 µM Plitidepsin, compared to controls (n=6 per condition per region). Histograms show means and error bars indicate standard deviation. P-values were defined using two-way ANOVA multiple comparisons (* P<0.05, ** P<0.01, *** P<0.001, and **** P<0.0001). Non-significant differences are not shown on the graph.

(B) Quantification of A549-ACE2 infected cells (%) at 24 hpi revealed by immunostaining using an anti-spike antibody at different multiplicities of infection (MOI).

(C) Cellular toxicity following Plitidepsin exposure at different concentrations, assessed via ATP assay at 24- and 48-hours posttreatment. The corresponding 50% cytotoxic concentrations (CC_50_) are indicated. Data are expressed as mean of mean ± Standard deviation of the mean, normalized to control.

(D) Percentage of SARS-CoV-2 infected cells (MOI= 0,1) at 24 hpi following cotreatments with Plitidepsin at different concentrations, measured by flow cytometry assay using an anti-SARS-CoV-2-spike antibody. The 50% inhibition concentration (IC_50_) is indicated. Results are represented as mean of mean ± standard deviation of the mean, normalized to control.

(E) Representative images of A549-ACE2 infected cells (MOI= 0,1) at 24 hpi, cotreated with 0, 0.01, 0.1, and 1 µM of Plitidepsin. Cells were stained with anti-spike antibody (red) and DAPI (blue).

(F) Representative images of lipid droplets in A549-ACE2 infected cells (MOI= 0,1) at 24 hpi compared to mock. Cells were cotreated with 0, 0.01, and 0.1 µM of Plitidepsin. Lipid droplets were stained with Bodipy^TM^ FLC12 (green). Scale bar represents 30 μm. **(F’)** Fluorescence intensity of Bodipy^TM^ FLC12 quantified and normalized to that of DAPI. P-values were defined using two-way ANOVA (* P<0.05, ** P<0.01, *** P<0.001, and **** P<0.0001).

