## Supplemental Figures S1-S7 for "Comprehensive investigation of SARS-CoV-2 intestinal pathogenesis in *Drosophila*"

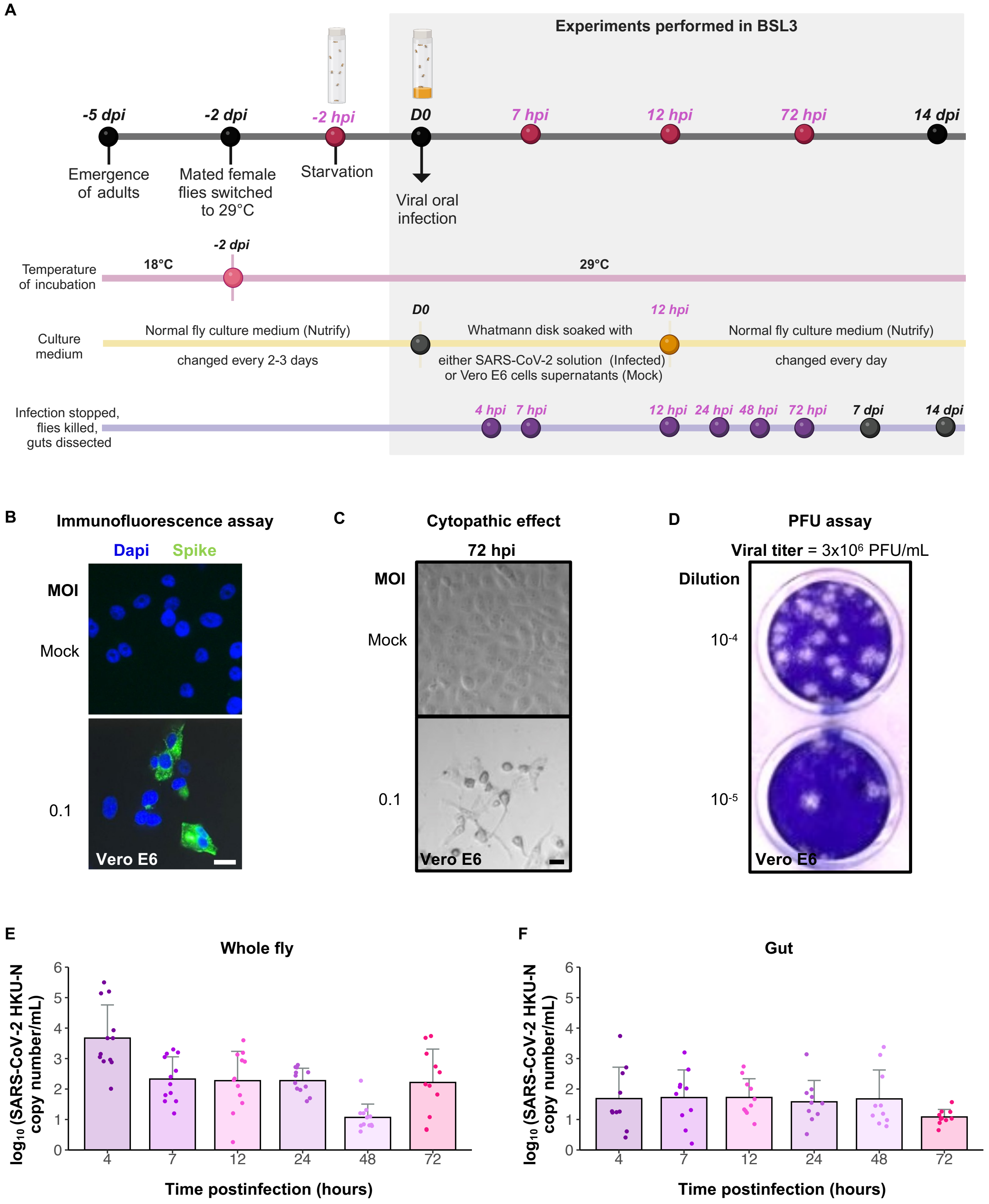

A
Experiments performed in BSL3
-5 dpi
-2 dpi
-2 hpi
D0
7 hpi
12 hpi
72 hpi
14 dpi
Emergence of adults
Mated female flies switched to 29°C
Starvation
Viral oral infection
Temperature of incubation
18°C
-2 dpi
29°C
Culture medium
Normal fly culture medium (Nutrify)
changed every 2-3 days
D0
Whatmann disk soaked with
either SARS-CoV-2 solution (Infected) or Vero E6 cells supernatants (Mock)
12 hpi
Normal fly culture medium (Nutrify)
changed every day
Infection stopped, flies killed, guts dissected
4 hpi
7 hpi
12 hpi
24 hpi
48 hpi
72 hpi
7 dpi
14 dpi
B
Immunofluorescence assay
Dapi
Spike
MOI
Mock
0.1
Vero E6
C
Cytopathic effect
72 hpi
MOI
Mock
0.1
Vero E6
D
PFU assay
Viral titer = 3x10⁶ PFU/mL
Dilution
10-4
10-5
Vero E6
E
Whole fly
log10 (SARS-CoV-2 HKU-N copy number/mL)
Time postinfection (hours)
4 7 12 24 48 72
0 1 2 3 4 5 6
F
Gut
log10 (SARS-CoV-2 HKU-N copy number/mL)
Time postinfection (hours)
4 7 12 24 48 72
0 1 2 3 4 5 6


**Figure S1**

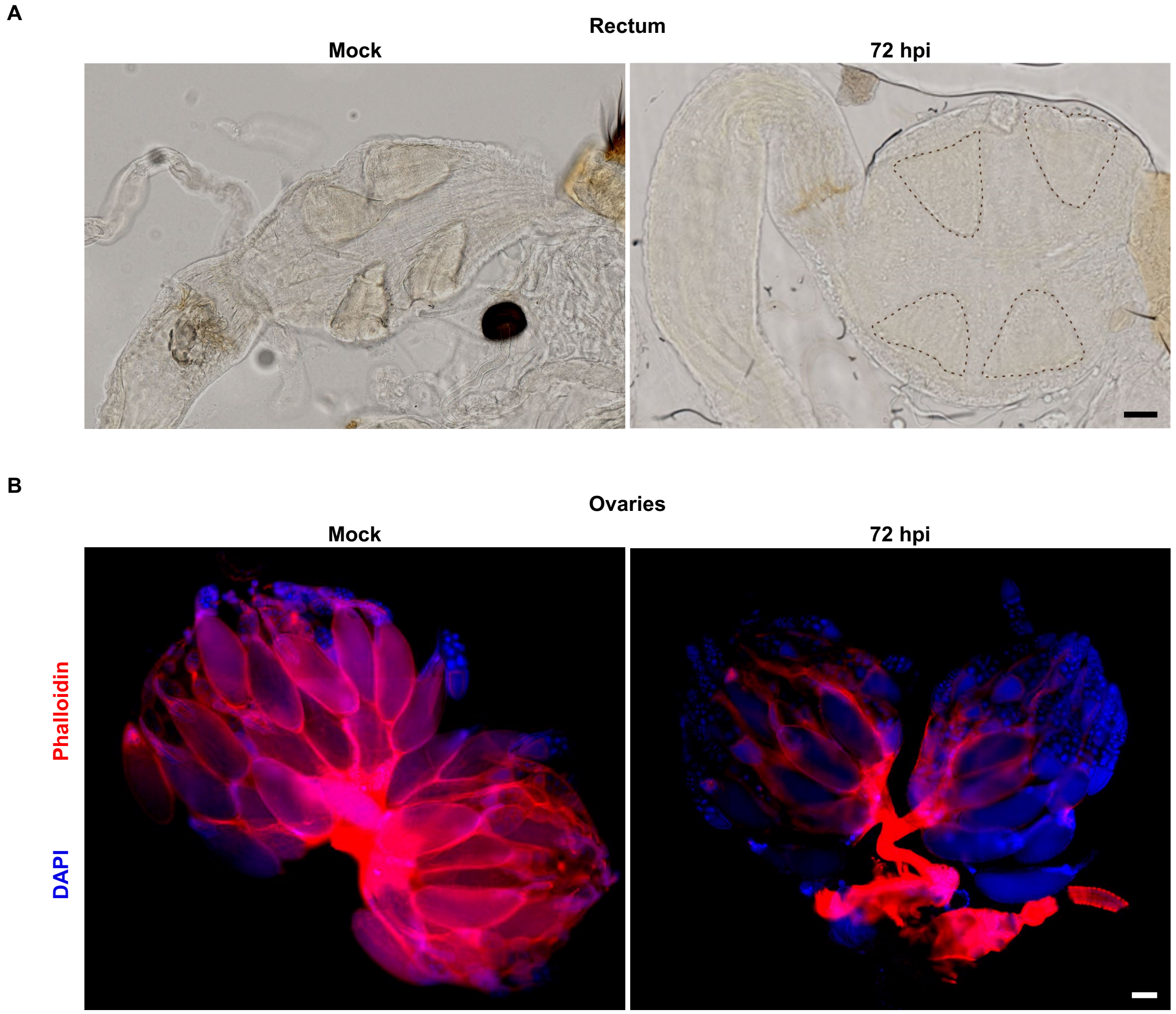


Figure S2

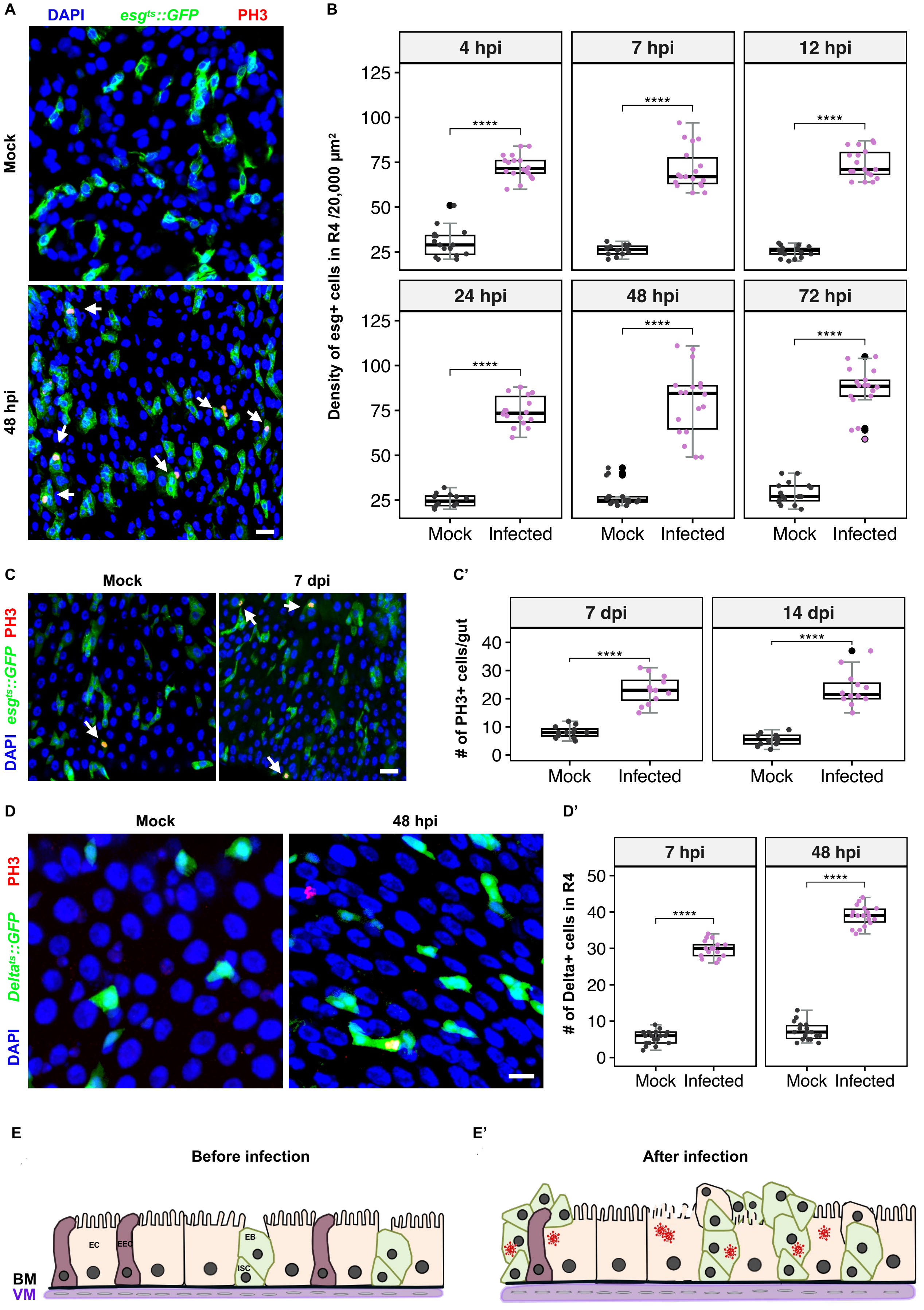
A
DAPI
esgts::GFP
PH3
Mock
48 hpi
B
4 hpi
7 hpi
12 hpi
24 hpi
48 hpi
72 hpi
Density of esg+ cells in R4 /20,000 µm²
Mock
Infected
C
Mock
7 dpi
DAPI esgts::GFP PH3
C'
7 dpi
14 dpi
### of PH3+ cells/gut
Mock
Infected
D
Mock
48 hpi
DAPI Deltats::GFP PH3
D'
7 hpi
48 hpi
### of Delta+ cells in R4
Mock
Infected
E
Before infection
EC
EEC
EB
ISC
BM
VM
E'
After infection

Figure S3

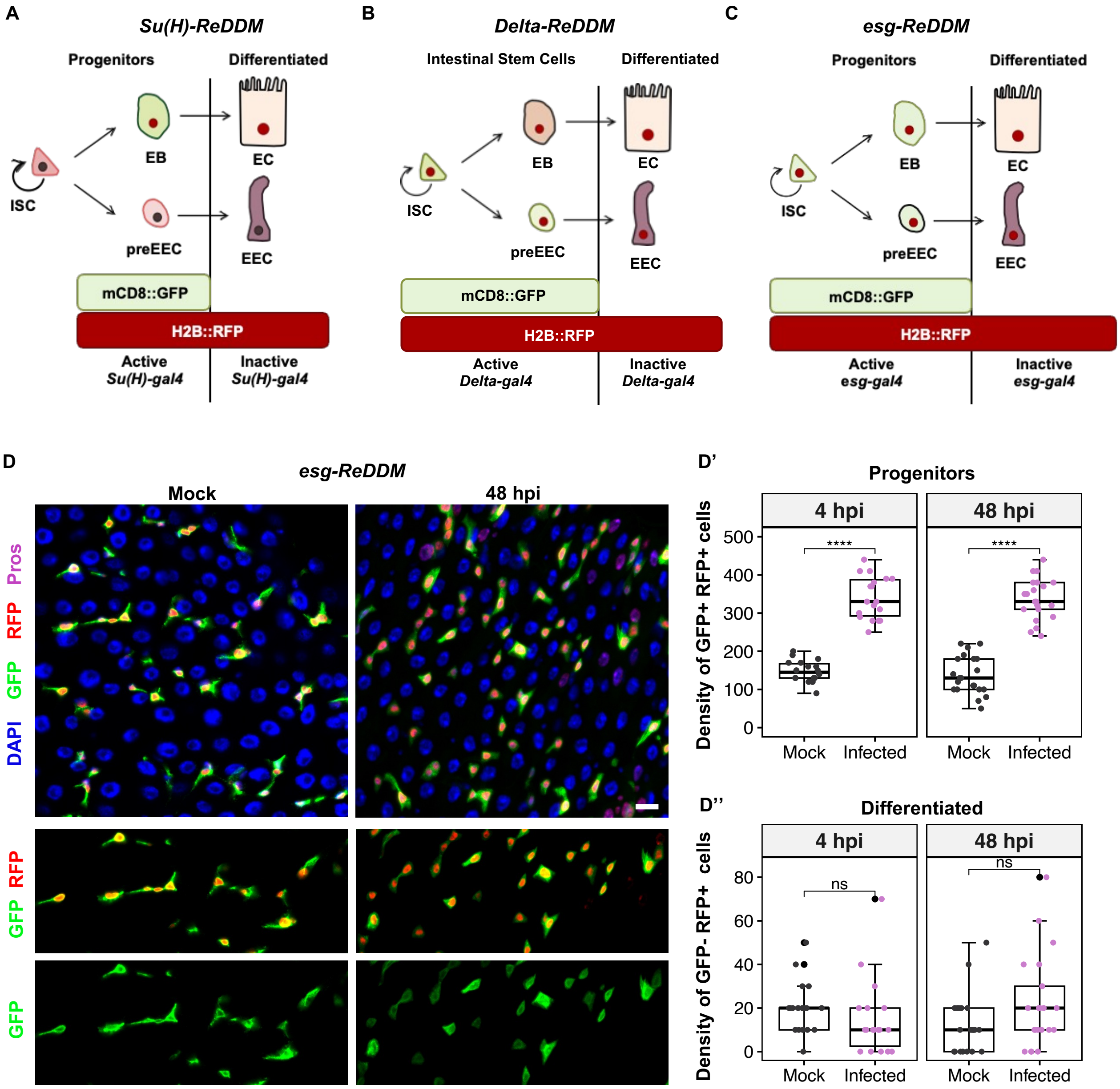
A
Su(H)-ReDDM
Progenitors
Differentiated
ISC
EB
EC
preEEC
EEC
mCD8::GFP
H2B::RFP
Active
Su(H)-gal4
Inactive
Su(H)-gal4
B
Delta-ReDDM
Intestinal Stem Cells
Differentiated
ISC
EB
EC
preEEC
EEC
mCD8::GFP
H2B::RFP
Active
Delta-gal4
Inactive
Delta-gal4
C
esg-ReDDM
Progenitors
Differentiated
ISC
EB
EC
preEEC
EEC
mCD8::GFP
H2B::RFP
Active
esg-gal4
Inactive
esg-gal4
D
esg-ReDDM
Mock
48 hpi
Pros
RFP
GFP
DAPI
RFP
GFP
GFP
D'
Progenitors
4 hpi
48 hpi
Density of GFP+ RFP+ cells
500
400
300
200
100
0
****
****
Mock
Infected
Mock
Infected
D''
Differentiated
4 hpi
48 hpi
Density of GFP- RFP+ cells
80
60
40
20
0
ns
ns
Mock
Infected
Mock
Infected

Figure S4

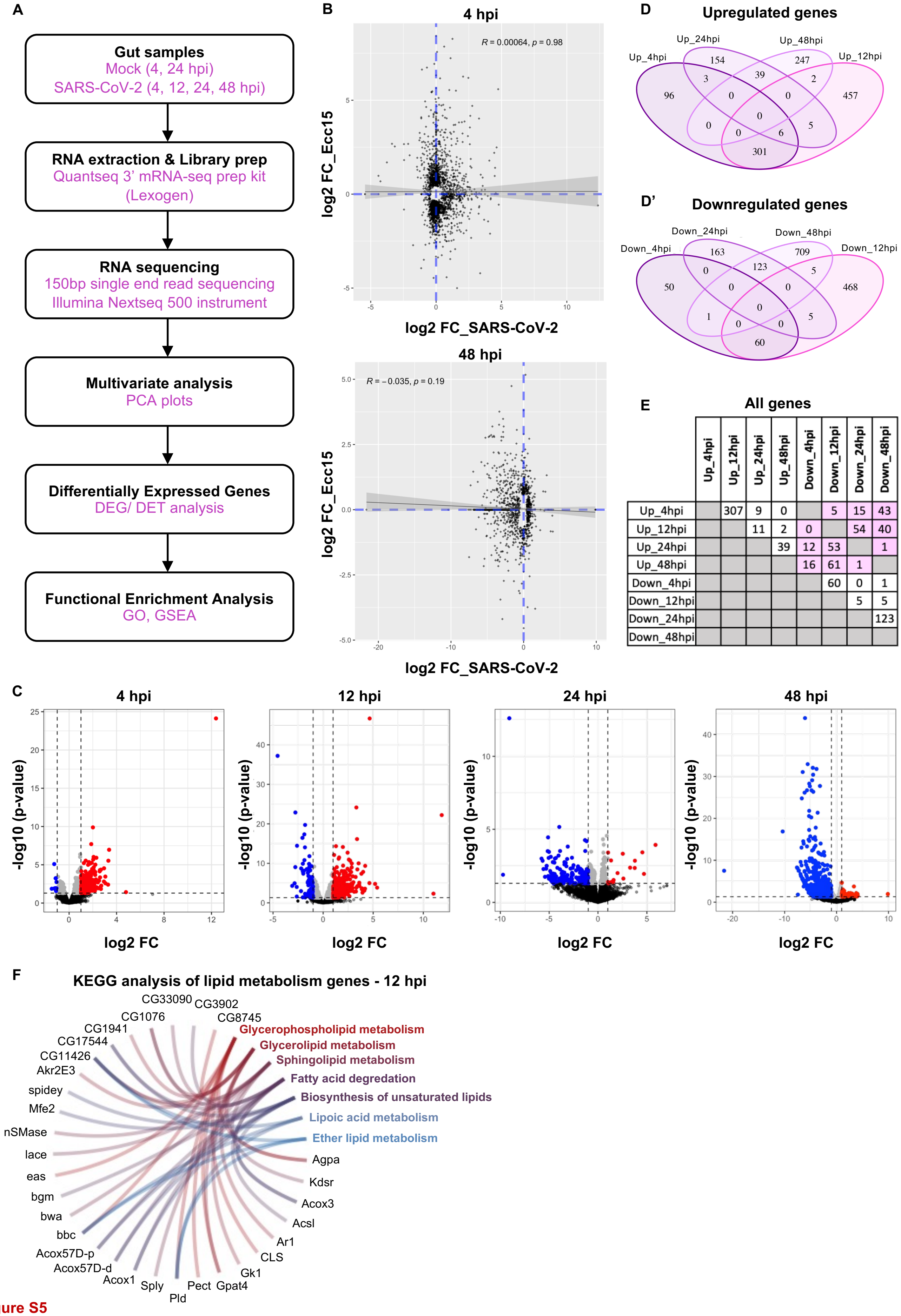


|  | Up_4hpi | Up_12hpi | Up_24hpi | Up_48hpi | Down_4hpi | Down_12hpi | Down_24hpi | Down_48hpi |
|---|---|---|---|---|---|---|---|---|
| Up_4hpi |  | 307 | 9 | 0 |  | 5 | 15 | 43 |
| Up_12hpi |  |  | 11 | 2 | 0 |  | 54 | 40 |
| Up_24hpi |  |  |  | 39 | 12 | 53 |  | 1 |
| Up_48hpi |  |  |  |  | 16 | 61 | 1 |  |
| Down_4hpi |  |  |  |  |  | 60 | 0 | 1 |
| Down_12hpi |  |  |  |  |  |  | 5 | 5 |
| Down_24hpi |  |  |  |  |  |  |  | 123 |
| Down_48hpi |  |  |  |  |  |  |  |  |

Figure S5

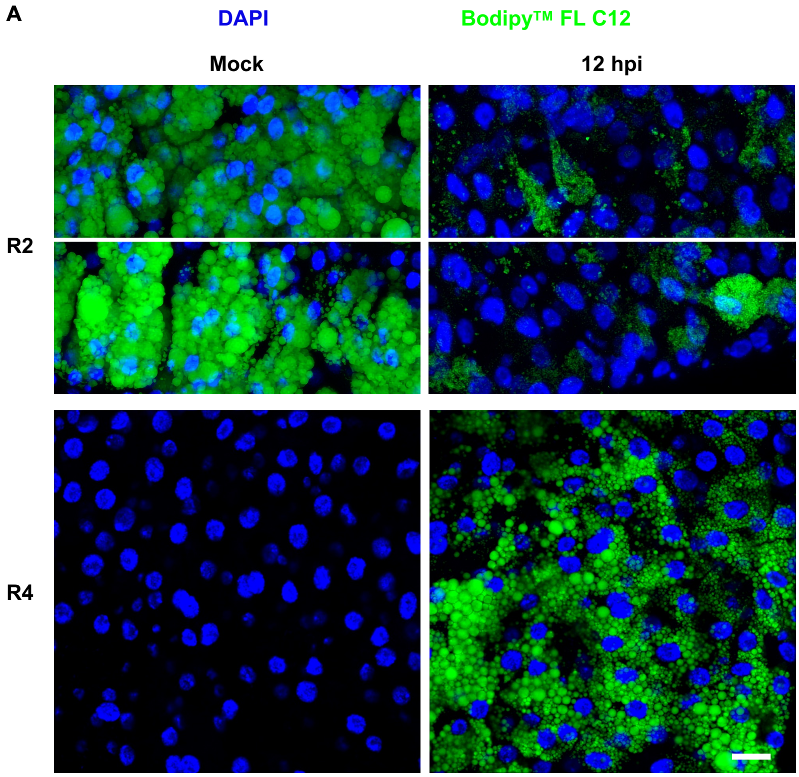


Figure S6

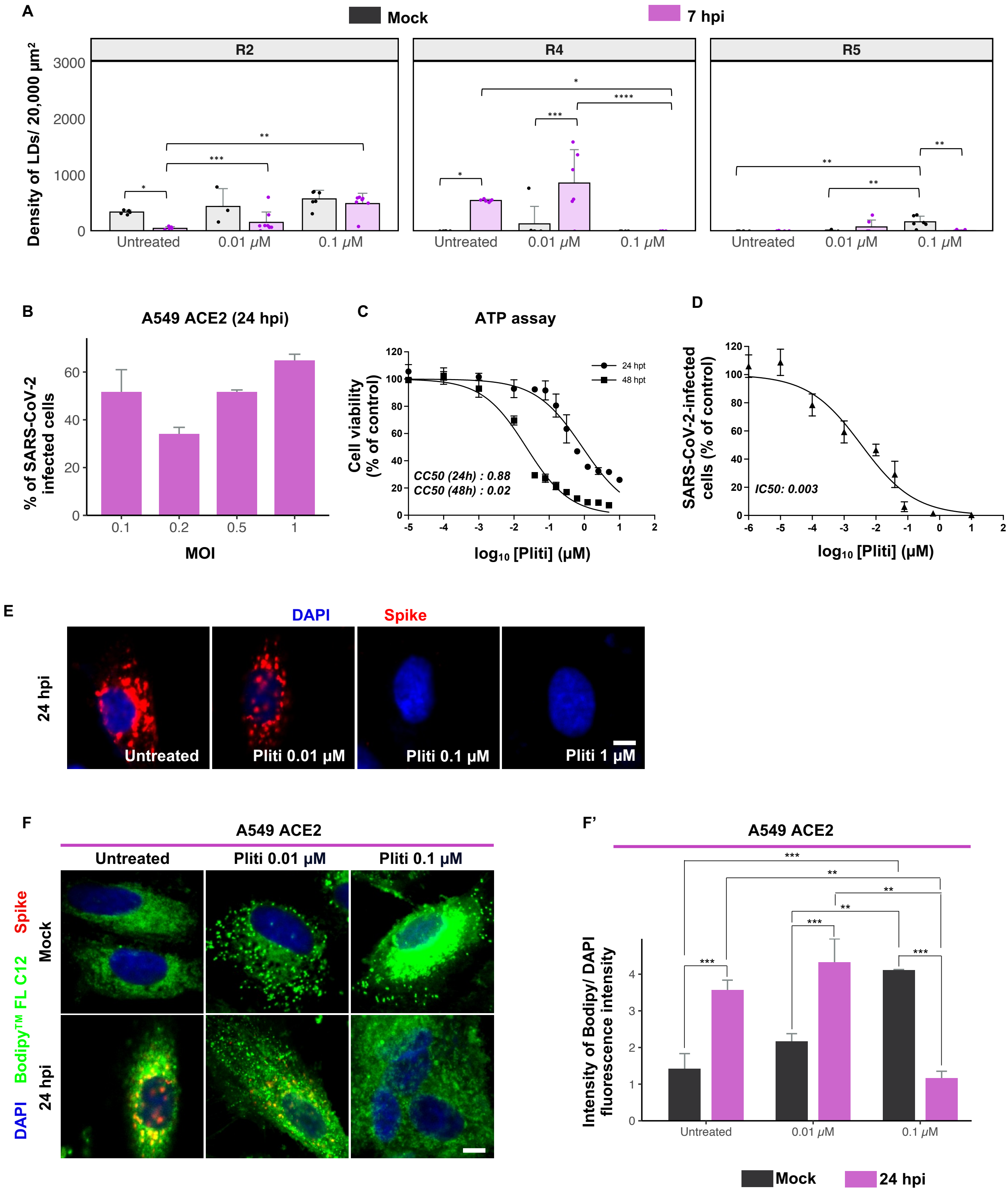

A
Mock
7 hpi
R2
R4
R5
Density of LDs/ 20,000 µm²
3000
2000
1000
0
Untreated
0.01 µM
0.1 µM
B
A549 ACE2 (24 hpi)
% of SARS-CoV-2 infected cells
MOI
0.1
0.2
0.5
1
C
ATP assay
Cell viability (% of control)
24 hpt
48 hpt
CC50 (24h) : 0.88
CC50 (48h) : 0.02
log10 [Pliti] (µM)
D
SARS-CoV-2-infected cells (% of control)
IC50: 0.003
log10 [Pliti] (µM)
E
DAPI
Spike
24 hpi
Untreated
Pliti 0.01 µM
Pliti 0.1 µM
Pliti 1 µM
F
A549 ACE2
Untreated
Pliti 0.01 µM
Pliti 0.1 µM
Mock
24 hpi
DAPI Bodipy™ FL C12 Spike
F'
A549 ACE2
Intensity of Bodipy/ DAPI fluorescence intensity
Untreated
0.01 µM
0.1 µM
Mock
24 hpi


Figure S7
